# Selective Disruption of *Plasmodium falciparum* mitochondrial DNA via G-Quadruplex-Binding Ligand RHPS4 Provides a Novel Antimalarial Strategy

**DOI:** 10.64898/2026.01.07.698092

**Authors:** Mariam Salim, Lucie Paloque, Thibaud Reyser, Flore Nardella, Jean-Michel Augereau, Yu Luo, Sébastien Britton, Jean-Louis Mergny, Virginie Gervais, Françoise Benoit-Vical, Dennis Gomez

## Abstract

Malaria caused by *Plasmodium falciparum* remains a major health threat, killing over 600,000 people annually. The spread of resistance to all major antimalarials, including artemisinins, highlights the urgent need for new drugs with distinct mechanisms of action. Here we show that the G-quadruplex ligand RHPS4, an acridine derivative, displays strong antiplasmodial activity against both drug-sensitive and -resistant *P. falciparum* strains and clinical isolates. RHPS4 primarily targets the trophozoite stage and induces major mitochondrial alterations, including reduction of mitochondrial DNA (mtDNA) and transcriptional dysfunctions. Bioinformatic analyses identified at least eight putative G4-forming sequences within the parasite’s mtDNA. Biophysical studies confirmed G4 folding of at least one sequence and its interaction with RHPS4. These findings indicate that RHPS4 disrupts *P. falciparum* mitochondrial metabolism through G4 stabilization, leading to parasite death, and establish mtDNA G4 structures as novel therapeutic targets for antimalarial development.

## Introduction

*P. falciparum* malaria still kills yearly around 600,000 people, the vast majority of whom being children under the age of 5 especially in sub-Saharan Africa (*1*). In recent decades, malaria treatment has relied on fewer than a dozen of antiplasmodial drugs, which mainly target heme polymerization (e.g., chloroquine), pyrimidine biosynthesis (e.g., sulfadoxine-pyrimethamine), cytochrome b activity (e.g., atovaquone), or lead to heme and proteins alkylation (e.g., artemisinin derivatives) (*2*). Unfortunately, *P. falciparum* has always been able to develop resistance to these drugs, including to the current major antimalarial drug, artemisinin (ART) and its derivatives (*3*, *4*). ART resistance is emerging and spreading in Africa (*5*) where 90% of malaria cases occur raising fears of a therapeutic dead end in the medium term. Now, the challenge is to develop drugs with novel modes of action to limit the risk of cross-resistance with existing drugs. Several new targets are under investigation, including ATPase 4, phosphatidylinositol-4-kinase, eukaryotic elongation factor 2, histone deacetylase 1, acetyl-CoA synthetase, and the proteasome (*6*). Guanine quadruplexes (G4, four-stranded non-canonical secondary structures of guanine-rich DNA or RNA sequences) (*7*, *8*) have been considered as potential targets to counteract parasite proliferation, and their selective stabilization by small chemical compounds, known as G4 ligands (*9*), has been shown to exert effects that, although modest, remain of significant interest (*10–15*). Over the past years, numerous studies have demonstrated the essential role of G4 on cell survival and cell metabolism regulation in different models (*16*, *17*). In cells, the stabilization of DNA-G4 structures by G4 ligands drives genomic instability, which is characterized by DNA breaks, DNA deletions, and chromosomal translocations (*18–24*). The formation of DNA-G4 structures is not restricted to the nuclear compartment. In human cells, G4 structures have been associated with mitochondrial DNA deletion breakpoints and with the stalling of the replication fork during mitochondrial DNA replication (*25*, *26*) contributing to regulation of mitochondrial gene expression and to perturbations of the mitochondrial genome (*27–29*). The presence of G4 structures in *Plasmodium* has been confirmed with specific G4-detecting antibodies (*13*, *15*). Given that *P. falciparum* genome is characterized by its extremely AT rich genome (> 80%) (*30*), only few regions of the parasite genome have the potential to form G4 structures (*10*, *13*, *14*, *31–33*). Interestingly, a genome-wide analysis of the nuclear and mitochondrial DNA of *Plasmodium* shows a higher density of G4 motifs in the mitochondrial DNA compared to the nuclear DNA (*31*).

To our knowledge, the impact of G4 ligands on *P. falciparum* mitochondrial genome has not yet been evaluated. Among several compounds already described as G4 ligands in mammalian cells, the acridine derivative RHPS4, was identified as a selective telomerase inhibitor through its capacity to bind and fold telomeric sequences into G4 (*34*). RHPS4 provokes telomeric dysfunctions and replication stress, induced by its capacity to impair fork progression in G4-containing regions in human transformed fibroblasts and melanoma cells (*35–37*). While most cellular studies performed with RHPS4 have focused on its telomeric effects, Falabella and co-workers reported that RHPS4 preferentially localizes into mitochondria where it abolishes mitochondrial functions and alters both mitochondrial replicative and transcriptional mechanisms (*28*).

In this report, we show that RHPS4 exhibits a substantial antiplasmodial activity against *P. falciparum* parasites, including on various drug-resistant laboratory strains and clinical isolates. We show that RHPS4 targets the trophozoite stage and induces important mitochondrial changes in *Plasmodium,* such as a strong reduction in mtDNA content coupled to transcriptional dysfunctions. Using bioinformatics analyses, we identify eight putative G4-forming sequences in the mitochondrial genome of the *P. falciparum* parasite and G4 folding of one sequence was experimentally confirmed. Furthermore, we demonstrated its interaction with RHPS4. Our study highlights RHPS4 as a promising antiplasmodial compound with a novel mechanism of action. These results open new avenues for antimalarial drug discovery targeting mitochondrial nucleic acid structures and underscore the potential of G4 ligands in combating drug-resistant malaria.

## RESULTS

### RHPS4 has a potent and selective antiplasmodial activity

The antiplasmodial activity of five chemically diverse G4 ligands (*52–56*) was assessed *in vitro* on different *P. falciparum* strains. Strikingly, only RHPS4 (Fig. 1A), with an IC_50_ value of 90 ± 10 nM, showed a strong antiplasmodial activity against the drug-sensitive strain F32-TEM (Table 1), ranging in the same order of magnitude that the three major antimalarial drugs ART, chloroquine (CQ) and atovaquone (ATQ).

**Figure 1:**
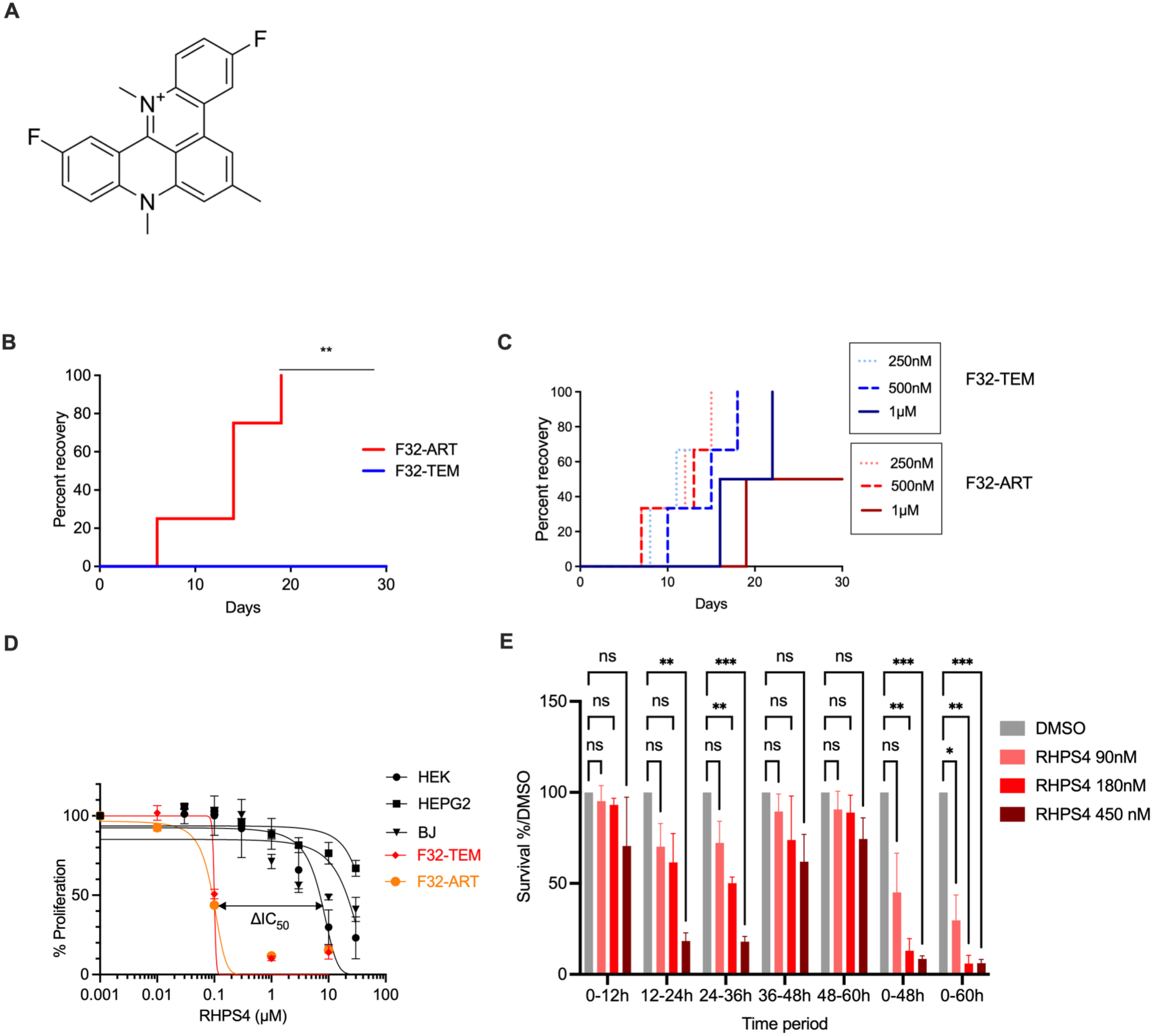
Selective antiplasmodial activity of the G-quadruplex ligand RHPS4. **A** Chemical structure of RHPS4**. B-C** Kaplan-Meier analyses of *P. falciparum in vitro* recrudescence tests comparing the ability of F32-ART (red color gradients) and F32-TEM (blue color gradients) strains to proliferate after treatments (ART 18 µM as control, RHPS4 250 nM, 500 nM and 1000 nM). The Mantel-Cox test was used for statistical analysis of the recrudescence data. (** = p<0.001). **D** Selectivity of RHPS4 toward parasites *versus* human cells. HEK, HepG2, and BJ-hTERT cell lines were treated for 72 h with a concentration range of RHPS4, and viability was measured by the sulforhodamine B assay. Selectivity index (SI) denotes the ratio between the CC₅₀ of the most sensitive human line (HEK) and the IC₅₀ of each *P. falciparum* strain. **E** Stage-specific activity of RHPS4. The F32-TEM strain was treated with RHPS4 using the IC₅₀ concentration, 2 × IC₅₀, or 5 × IC₅₀, initiated at successive 12-h intervals during five successive 12-h periods from a highly-synchronized ring stage population. Following each pulse, parasites were washed and cultured drug-free until 60 h. The % of survival was then determined using stained blood smear and normalized to DMSO treatment. Error bars indicate mean ± SD (n = 3 independent experiments).

**Table 1.**
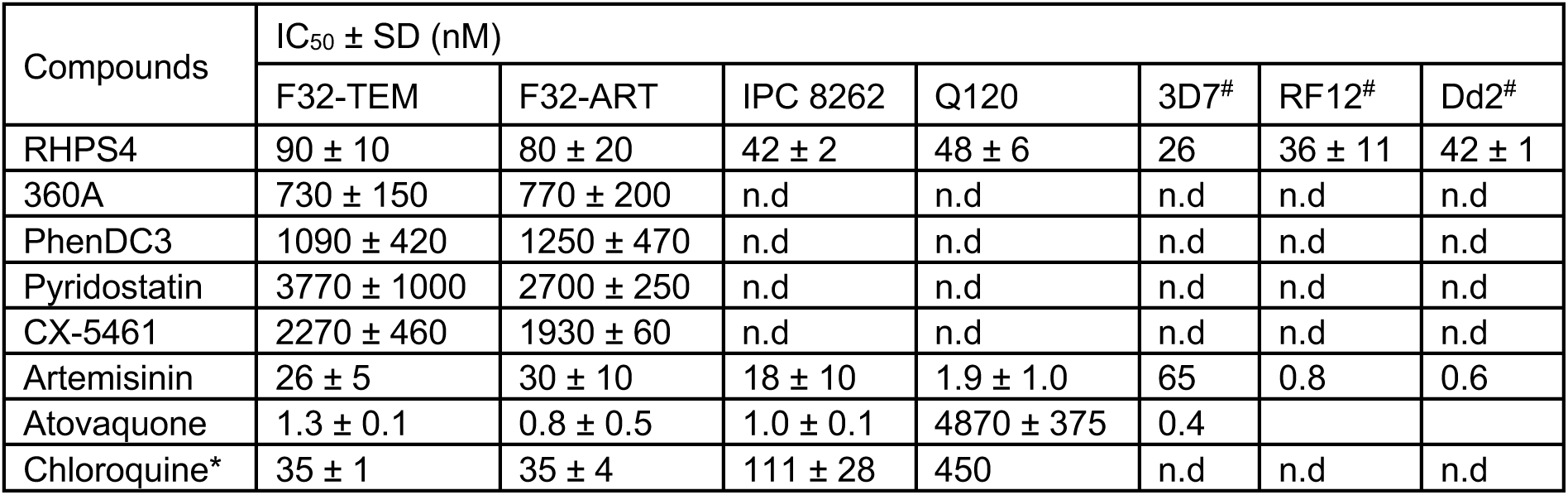
IC₅₀ values of G-quadruplex ligands and reference antiplasmodial drugs against *Plasmodium falciparum* strains. IC₅₀ values of various G4 ligands, and reference antiplasmodial drugs (ART: artemisinin, ATQ: atovaquone and CQ: chloroquine) on several *Plasmodium falciparum* strains: F32-TEM (drug-sensitive), F32-ART (ART-resistant), IPC8262 (resistant to both ART and CQ), and Q120 (resistant to both ATQ and CQ). IC₅₀ values were determined based on DNA quantification after 48 hours of treatment with a concentration range for each compound. Data represent the mean of five independent experiments (n= 5). For 3D7 (drug sensitive, n = 1), RF12 (CQ-resistant and ART-resistant, n = 2) and Dd2 (CQ-resistant, n = 2) strains, the data were obtained by Medicine for Malaria Venture using the LDH assay and 72h incubation. *\** These values have been previously published in (*57*). *^#^* data from MMV using Pf LDH assay and 72h incubation.

| Compounds | IC<sub>50</sub> ± SD (nM) |  |  |  |  |  |  |
| --- | --- | --- | --- | --- | --- | --- | --- |
|  | F32-TEM | F32-ART | IPC 8262 | Q120 | 3D7<sup>#</sup> | RF12<sup>#</sup> | Dd2<sup>#</sup> |
| RHPS4 | 90 ± 10 | 80 ± 20 | 42 ± 2 | 48 ± 6 | 26 | 36 ± 11 | 42 ± 1 |
| 360A | 730 ± 150 | 770 ± 200 | n.d | n.d | n.d | n.d | n.d |
| PhenDC3 | 1090 ± 420 | 1250 ± 470 | n.d | n.d | n.d | n.d | n.d |
| Pyridostatin | 3770 ± 1000 | 2700 ± 250 | n.d | n.d | n.d | n.d | n.d |
| CX-5461 | 2270 ± 460 | 1930 ± 60 | n.d | n.d | n.d | n.d | n.d |
| Artemisinin | 26 ± 5 | 30 ± 10 | 18 ± 10 | 1.9 ± 1.0 | 65 | 0.8 | 0.6 |
| Atovaquone | 1.3 ± 0.1 | 0.8 ± 0.5 | 1.0 ± 0.1 | 4870 ± 375 | 0.4 |  |  |
| Chloroquine* | 35 ± 1 | 35 ± 4 | 111 ± 28 | 450 | n.d | n.d | n.d |

We next evaluated all compounds against the ART-resistant strain F32-ART (*51*). As previously observed, most of G4 ligands remained weakly active, while RHPS4 maintained a strong antiproliferative effect against the F32-ART strain, with an IC_50_ value of 80 ± 20 nM. This activity was also similar in the multidrug-resistant clinical field isolates IPC8262 (resistant to ART and CQ), and Q120 (resistant to ATQ and CQ) with IC_50_ values of 42 ± 2 nM and 48 ± 6 nM, respectively, showing no cross-resistance between RHPS4, CQ and ATQ (Table 1). These results were confirmed in another laboratory with a different assay against three other strains (3D7, RF12 and Dd2), as part of a collaboration with Medicines for Malaria Venture (MMV) (Table 1). Due to the quiescence-based mechanism of ART resistance (*51*), IC_50_ determination cannot provide evidence of cross-resistance with ART, which is assessed through a recrudescence assay based on the difference between ART-resistant and ART-sensitive parasites in their ability to proliferate after treatment (*43*). As expected, when treated with ART 18 μM for 48 h, the ART-resistant F32-ART strain was able to resume its growth after drug washout (median recovery time of 14 days), while no ART-sensitive parasites survived (no recrudescence of F32-TEM at day 30) (Fig. 1B). Using the same assay, the recrudescence capacities of the two strains F32-TEM and F32-ART were assessed after treatment with 250 nM, 500 nM and 1 µM of RHPS4 (corresponding to three, six, and 12 times the IC_50_ value of RHPS4).

While a short delay in recrudescence was observed with 1 µM of RHPS4, the recrudescence capacities of the two strains F32-TEM and F32-ART were not statistically different at all concentrations (*p-*value = 0.69, 0.78 and 0.43 for 250 nM, 500 nM and 1 µM respectively), demonstrating the absence of cross-resistance between ART and RHPS4 (Fig. 1C). An important point for antimalarial drugs profile is their selective activity against parasites versus the host cells. This was evaluated using cytotoxicity assays on three different human cell lines: HepG2 (derived from hepatocarcinoma), HEK (derived from embryonic kidney) and BJ-hTERT cells (immortalized non-transformed human fibroblast). CC_50_ values of RHPS4 towards human cells were close to or higher than 10 µM (Fig. 1D), indicating a highly selective action of RHPS4 against *P. falciparum* parasites compared to human cells with a selectivity index (SI) >100.

### RHPS4 is active during the trophozoite stage

During its 48 h erythrocytic cycle, *P. falciparum* matures and multiplies through different stages: ring, trophozoite and schizont. This ultimately leads to the release of new parasites into the bloodstream, initiating a new cycle. To evaluate the stage-specificity of RHPS4 action, we determined its activity at different concentrations during five successive 12 h periods from a highly-synchronized ring-stage population. As a control, we treated the parasites for 48 h and 60 h. Consistent with our IC_50_ results, a 50% inhibitory effect was observed after 48 h of treatment with 90 nM of RHPS4 (IC_50_ value). The parasite survival was affected by increasing both the concentration and the incubation time. The impact of RHPS4 on *P. falciparum* proliferation is markedly stage-dependent. Indeed, a minor impact on parasitemia was noted for 12 h treatments initiated at 0, 36, and 48 h, even at the highest RHPS4 concentration. In contrast, a pronounced effect on survival was observed when the 12 h treatment was initiated at 12 or 24 h, 450 nM of RHPS4 inducing the most drastic effect (Fig. 1E, supplementary Fig. S1).

Overall, our results indicate that RHPS4 acts mainly during the 12–36 h period of the parasite asexual blood cycle, which coincides with the trophozoite stage.

### RHPS4 is slow-acting against *P. falciparum* parasites

The speed of action of a compound determines the exposure time required to achieve antiplasmodial activity. We determined the parasite reduction ratio (PRR) of *P. falciparum* F32-TEM strain in response to treatment with 10 times the IC_50_ of RHPS4 for up to 120 h, using ATQ and ART as controls. RHPS4 showed a slow-acting parasite reduction profile similar to ATQ with a time to achieve the maximum speed of action (lag phase) of 48 h, a parasite reduction ratio of 3.6 log over 48 h (Log(PRR)) and a 99.9% parasite clearance time (PCT) of 70 h (Fig. 2A). On the contrary, the fast-acting compound ART presented a 99.9% PCT value of 16 h with no lag phase and a Log(PRR) value greater than 5 (extrapolated from the first 24 h of treatment) (Fig. 2A). These data were confirmed by monitoring the parasite morphology throughout the treatment period (Supplementary Fig. S2A). In control condition (DMSO), reinvasion was observed every 48 h (full cycle), as evidenced by the rings observed corresponding to the beginning of a second cycle. In contrast, after 72 h of continuous exposure to RHPS4 or ATQ, intra-erythrocytic development was blocked at the trophozoite stage. These microscopic observations indicate that RHPS4 and ATQ achieve antiplasmodial activity at the trophozoite stage and confirm the PRR data with a lag phase of 48 h. The rapid onset of action of ART is also confirmed by the fact that, after 24 h of continuous exposure to artemisinin, parasites are dead with only pyknotic parasites visible in the smear.

**Figure 2:**
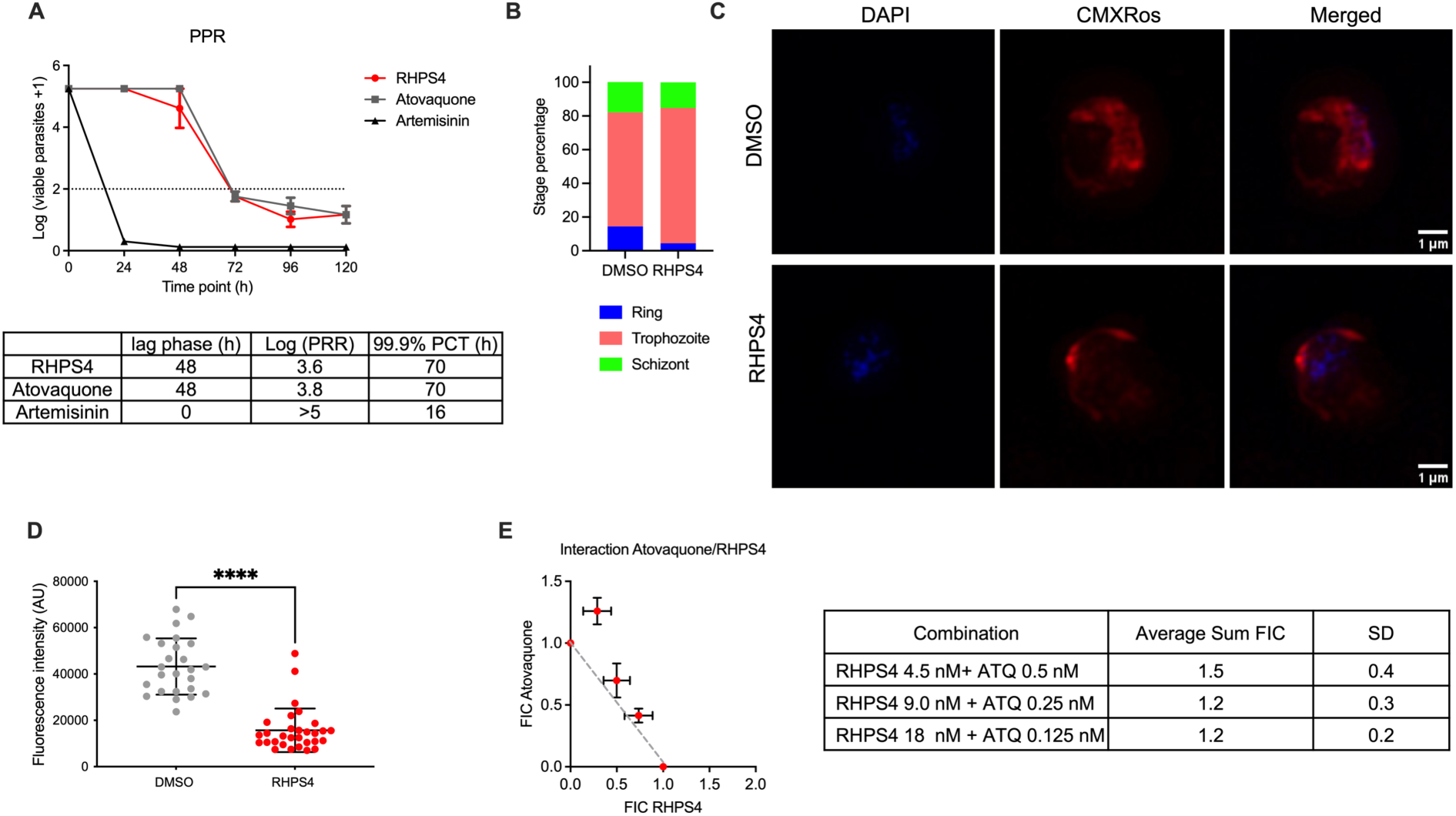
Mitochondrial effects, and drug interaction profile of RHPS4 in *Plasmodium falciparum*. **A** Determination of the parasite reduction ratio (PRR): *in vitro* lag phase, parasite reduction ratio and 99.9% parasite clearance times (PCT) in response to RHPS4, atovaquone and artemisinin exposures at a concentration corresponding to 10 × IC_50_. Lag phase: time needed to achieve the maximum rate of killing. Log (PRR): log10 of parasite reduction ratio over one cycle (48h) when maximum speed of action is achieved. 99.9% PCT: time required to reduce 99.9% of the initial parasite population. **B-D** Microscopic evaluation of mitochondrial activity in F32-TEM trophozoites (highly synchronized at 24-28 h post-invasion) after a 6 h treatment with either DMSO or 180 nM RHPS4: (B) Percentage of each parasite stages (ring, trophozoite and schizont) counted by two independent experienced microscopists on stained blood smears. Morphologically, ring-stage parasites were defined as parasites with a ring shape, trophozoite stages as parasites with full cytoplasm, and schizonts as parasites with punctate nuclear coloration of the developing merozoites. (C) Super resolution fluorescence images of parasites stained with MitoTracker Red CMXRos (1 h at 500 nM) to visualize mitochondria and with DAPI (2 µg/mL, 5 min) to visualize nuclei. Images were acquired on a Zeiss Elyra super resolution microscope. (D) Mitochondrial fluorescence was quantified with ImageJ software by measuring the mean MitoTracker intensity per infected RBC. Bars show mean ± SD from 26 infected red blood cells for DMSO and 30 infected red blood cells for RHPS4; *p-*values were calculated with a two tailed Student’s t test. **E** Isobologram and sum of fractional IC_50_ (FIC) of the interaction between RHPS4 and ATQ on *P. falciparum* F32-TEM strain. Error bars represent the standard deviation of four independent experiments. In the case of an additive combination of the two molecules, the isobologram lies on a straight line. In the case of a synergistic combination, the isobologram lies below the line, while in the case of antagonistic combinations, it lies above it. The FIC is the ratio of the IC_50_ of the combination to the IC_50_ of the molecule alone. More precisely, ΣFICs <1 denote synergism, ΣFICs ≥1 and <2 denote additive interaction, ΣFICs ≥2 denote antagonism, and ΣFICs ≥4 denote marked antagonism (*59*).

These results show that RHPS4 is a slow-acting compound, similarly to ATQ, a well-known inhibitor of *P. falciparum* mitochondrial metabolism targeting cytochrome b (*58*).

### RHPS4 impacts *P. falciparum* mitochondrial metabolism

Initially described as a selective telomerase inhibitor and shown to alter telomeric functions, RHPS4 has also been reported to impair mitochondrial metabolism in mammalian cells (*28*). Mitochondrial dysfunctions induced by RHPS4 appear at low concentrations (nanomolar range), with no impact on nuclear DNA, indicating a preferential accumulation in the mitochondrial organelle (*28*). Based on these data, we evaluated the impact of RHPS4 on *P. falciparum* mitochondrial metabolism. Treatment of highly synchronized trophozoites (24-28 h post-invasion, corresponding to the most susceptible stage, Fig. 1E) with 180 nM RHPS4 for 6 h did not impact the parasite distribution in the different stages analyzed using stained thin blood smears (Fig. 2B).

In parallel, we assessed *Plasmodium* mitochondrial activity using MitoTracker Red CMXRos staining, which depends on the mitochondria membrane potential and redox state. Microscopic analysis of the fluorescence intensity of *Plasmodium* in infected red blood cells (DAPI + / CMXRos +) showed a significant decrease in mitochondrial staining for RHPS4-treated parasites, which was reduced by three folds compared to the control (*p*-value < 0.0001), indicative of an altered mitochondrial membrane potential (Fig. 2C-D and Supplementary Fig. S2B-C). Taken together, these results indicate that a short RHPS4 treatment leads to a mitochondrial dysfunction at the trophozoite stage without altering parasite morphology and/or cycle progression.

### Additive effect of RHPS4 with atovaquone

As RHPS4 seems to affect mitochondria, we studied its interaction with ATQ. As a control, we tested the combination ATQ + proguanil as this synergistic combination is used in therapy. As expected, we obtained a synergistic effect with sums of fractional IC_50_ values below 1, consistent with a previous study (Supplementary Fig. S2D)(*60*). By contrast, the combination of RHPS4 with atovaquone shows an additive effect as the sums of the fractional IC_50_ values are close to 1 (Fig. 2E).

### RHPS4 alters nucleic acid transactions in *Plasmodium* mitochondria

Because mitochondrial dysfunctions induced by RHPS4 in mammalian cells are associated with alterations in nucleic acid transactions, qPCR assays were performed to evaluate RHPS4 impact on F32-TEM mtDNA. The 6 kb *P. falciparum* mitochondrial genome encodes for several small ribosomal RNA subunits and for three essential components of the ATPase complex: cytochrome c oxidase subunit 1 (COX1), cytochrome c oxidase subunit III (COX3), and cytochrome b (CYTB). We extracted the total parasite genomic DNA after 48 h of exposure to 90 nM RHPS4 (corresponding to the IC_50_ value), starting with trophozoite parasites, and quantified the relative abundance of mtDNA by qPCR with primers targeting different mtDNA regions (coding for ribosomal RNA fragments RNA1 and RNA4 and COX1, COX3 and CYTB coding regions). Treatment with RHPS4 resulted in five to ten-fold reduction in mtDNA relative abundance compared with the DMSO control for all DNA regions tested (Fig. 3A and Supplementary Fig. S3A). As controls, mtDNA abundance was evaluated on parasites treated with the two antimalarial drugs ART and ATQ. After 48 h treatment at their respective IC_50_ concentrations, no significant reduction was observed in mtDNA relative abundance compared with the DMSO condition (Fig. 3A). Importantly, RHPS4 did not interfere with the PCR assay itself since the addition of up to 10 µM RHPS4 in the PCR mix did not impact, *in vitro,* mtDNA amplification (Supplementary Fig. S3B). These results strongly suggests that RHPS4 operates through an original mechanism directly related to mtDNA, as compared to antimalarial drugs. A similar result, with a five-fold reduction in mtDNA abundance after RHPS4 treatment, was obtained with the field isolates Q120 and IPC8262 (Fig. 3B), indicating a shared mechanism of action of RHPS4 independent of the *P. falciparum* strain. Finally, we observed the same significant effect on mtDNA abundance when highly-synchronized parasites (12-16h post-invasion) were treated for 24 h with 180 nM of RHPS4, to cover the trophozoite stage, showing that RHPS4-induced mtDNA depletion correlated with its stage-specific activity (Fig. 3C). Based on these results, we hypothesized that the production of mitochondrial mRNAs and rRNAs should be equally affected by the RHPS4 treatment. RT-qPCR analyses were performed on the total RNA extracted from highly-synchronized trophozoites treated with 180 nM RHPS4 for 24 h. RT-qPCR analyses showed a five to ten-fold reduction of all tested transcript levels in treated parasites relative to control, which is consistent with the depletion of mtDNA induced by RHPS4 (Fig. 3D), indicating a global impact of RHPS4 on *P. falciparum* mtDNA transcription. As shown in Fig. 1B, RHPS4 exhibits a marked antiproliferative effect compared to other G-quadruplex ligands. To determine whether there is a correlation between the antiproliferative effects of these compounds and their ability to affect the mitochondrial genome of the parasite, comparative qPCR analyses were performed. As shown in Figure 3E, Pyridostatin and CX-5461 had no significant effect on mtDNA abundance following treatment at concentrations corresponding to their respective IC₅₀ values. In contrast, under the same treatment conditions, PhenDC3 and 360A—both charged compounds—induced significant reductions in mtDNA content, by approximately 50% and 65% respectively, relative to the DMSO control. However, the reduction of mtDNA by these two compounds was significantly lower than the massive reduction achieved by RHPS4 (Supplementary Fig. S3C). Altogether, these data suggest a positive correlation between the ability of different ligands to perturb the mitochondrial genome of the parasite and their antiproliferative activity.

**Figure 3:**
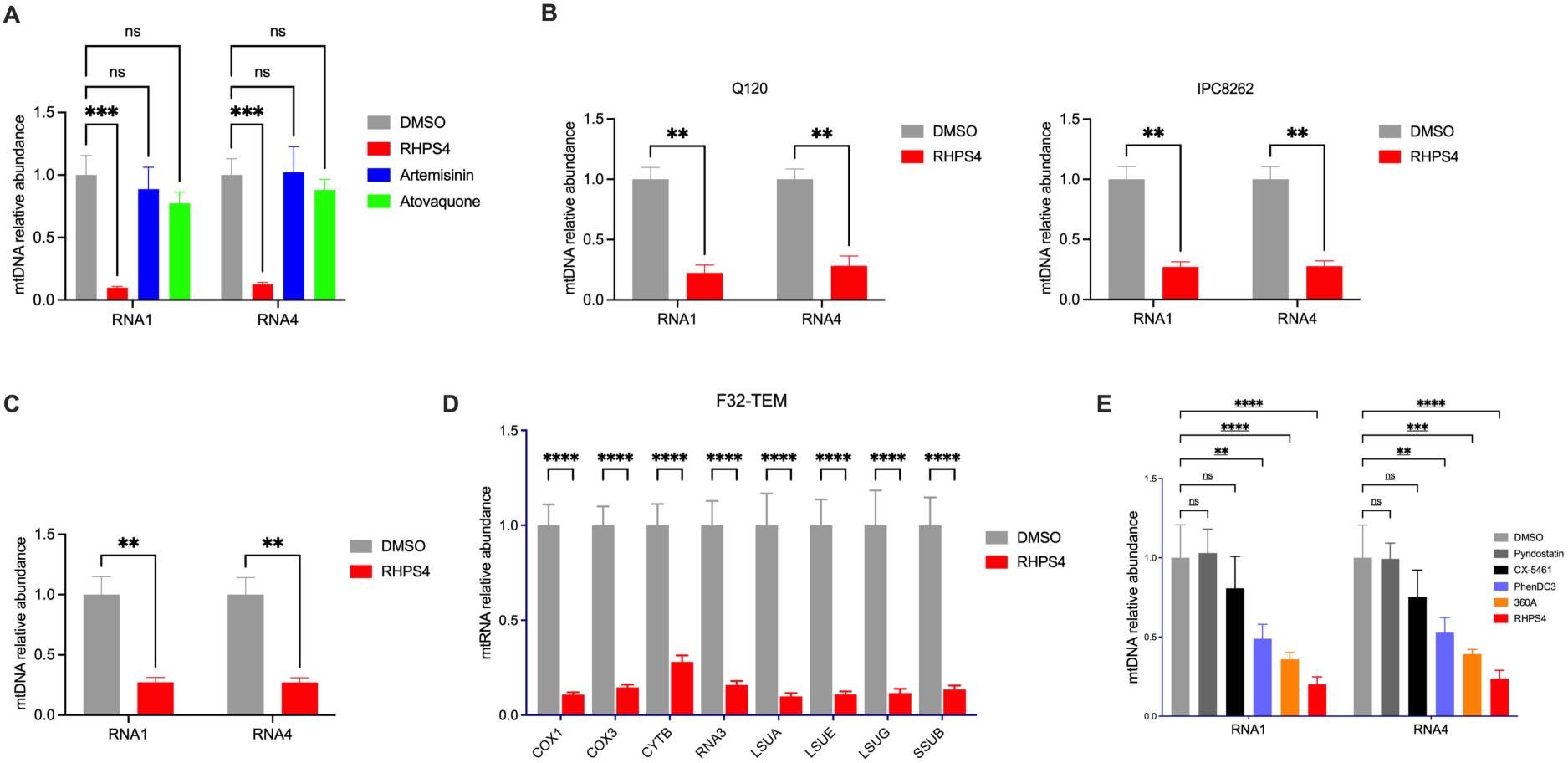
RHPS4 alters mitochondrial DNA abundance in *Plasmodium falciparum* and clinical isolates. **A-C** Relative abundance of different mtDNA sequences at the trophozoite stage, measured by qPCR after a 48-h exposure to RHPS4, artemisinin, and atovaquone at their respective IC₅₀ values (90, 20 and 5 nM, respectively) using the F32-TEM strain (A), Q120 (B), and IPC8262 (C). For both clinical isolates, which present a slightly slower replicative cycle than laboratory strains, data were obtained after a 72-h treatment. Bars show mean ± SD from n = 3 independent experiments for F32-TEM and from n = 2 independent experiments for both clinical isolates. **D** Relative abundance of mtDNA sequences measured by qPCR after a 24 h exposure to 180 nM RHPS4 of highly-synchronized F32-TEM parasites (12-16h post-invasion). The mtDNA abundance was normalized to the nuclear sequence PF3D7_1371000 coding for ribosomal RNA 18S. **E** Relative abundance of mtDNA sequences measured by qPCR after 48 h of exposure to different G4 ligands at their respective IC₅₀ concentrations. mtDNA abundance was normalized to the nuclear gene PF3D7_1371000, which encodes 18S ribosomal RNA. Bars show mean ± SD from n = 3 independent experiments.

### RHPS4 interacts with a G-quadruplex forming sequence from *P. falciparum* mtDNA

*In silico* analysis of *P. falciparum* mtDNA indicated the presence of at least eight sequences having the potential to form G-tetrad G4 structures (Supplementary Table 2) (in order not to miss any G4 candidate, relatively permissive parameters were chosen to identify these motifs). Two of them, Pf_mtDNA_G4_1 and Pf_mtDNA_G4_4, were predicted to fold into a G4 by three different algorithms (Table 2).

**Table 2:**
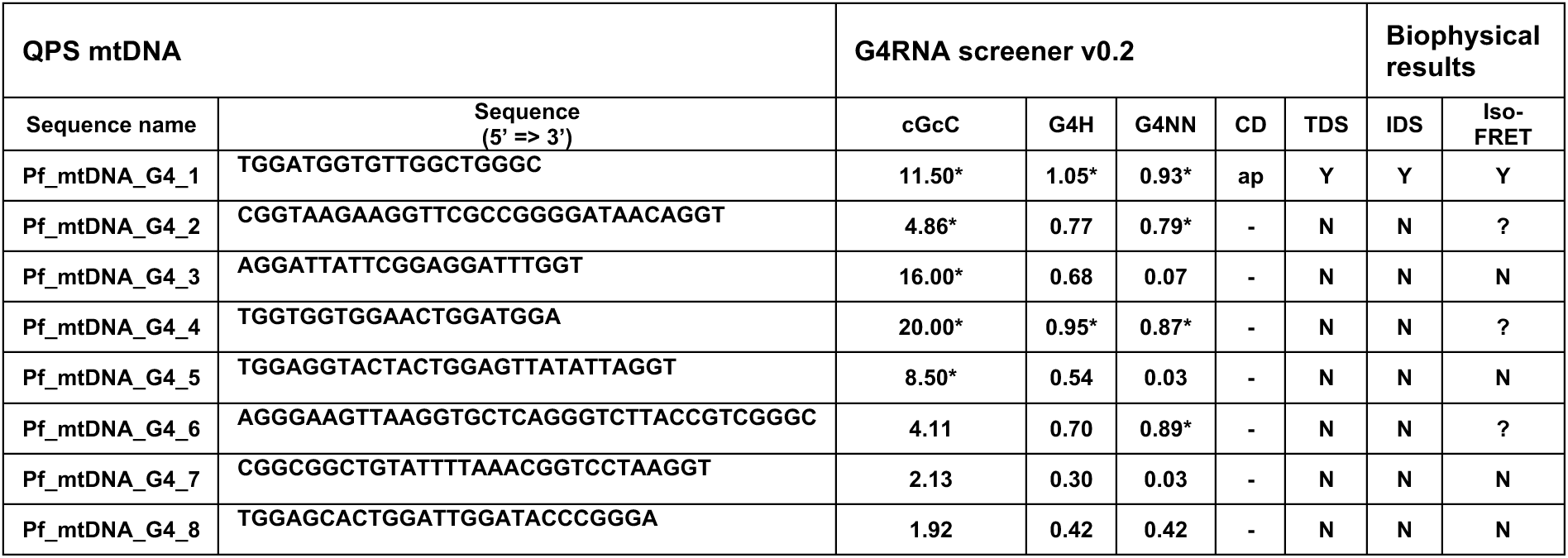
mtDNA sequence choice and summary of biophysical results. Eight candidate DNA sequences were selected based on scores according to three different algorithms (cGcC, G4H for G4Hunter and G4NN). Scores above arbitrary thresholds are considered to be potential candidates and are indicated with a “*”. The biophysical results are summarized in the last four columns. For the CD spectra, “ap” stands for a antiparallel G4 structure; “-“: inconclusive. For TDS, IDS and iso-FRET, “Y”, “N” and “?” stand for “Yes”, “No” and “unconclusive”, respectively (whether the results are similar to what is observed for known G4-forming sequences).

In order to assess their folding *in vitro*, short DNA oligonucleotides were analyzed through biophysical methods. Eight sequences were first characterized using circular dichroism (CD). CD is a powerful and standard method for evaluating the folding properties of nucleic acids and characterizing G4 topologies (parallel-stranded and antiparallel-stranded G-tetrad cores) through distinct CD signatures. In the K^+^ conditions used, one of the eight sequences, Pf_mtDNA_G4_1, exhibited a CD profile with a positive band at ≈ 295 nm and a negative signal at ≈ 260 nm, a profile similar to those encountered for anti-parallel G4s (Fig. 4A). To further characterize these sequences, UV spectroscopy analyses and fluorescence-based assays were performed. Isothermal differential spectra (IDS) and Thermal differential spectra (TDS) are two simple spectroscopic methods to demonstrate G4 formation based on the comparison of absorbance spectra of the folded and unfolded forms (*47*, *48*). These methods involve comparing spectra recorded under permissive conditions (K^+^ for IDS, 25°C for TDS) for G4 formation and non-permissive conditions (Li^+^ for IDS, 95°C for TDS). Under these conditions IDS and TDS showed no significant absorbance changes at 295 nm for 7 sequences out of the 8 considered. Again, Pf_mtDNA_G4_1 sequence was an exception, with TDS and IDS spectra reminiscent of G4-forming sequences, in agreement with its folding into a G4 *in vitro* (Supplementary Fig. S4A-B). Finally, iso-FRET was used to demonstrate G4 formation. Iso-FRET is an isothermal fluorescence-based method which evaluates the ability of DNA oligonucleotides to bind a highly selective G4 ligand, PhenDC3, under competitive conditions with a well-characterized G4-forming sequence (*49*). In this assay, a reduction in the fluorescence signal resulting from the displacement of PhenDC3 from the reporter sequence indicates the propensity of the tested sequence to fold into a G4 structure *in vitro* and act as a competitor for PhenDC3. As shown in Supplementary Fig. S4C, Pf_mtDNA_G4_1 induced a significant reduction in fluorescence emission as compared to other sequences tested, consistent with its propensity to adopt a G4 structure *in vitro*. Overall, four independent methods (CD, TDS, IDS, iso-FRET) demonstrated the ability of Pf_mtDNA_G4_1 to adopt a quadruplex fold.

**Figure 4:**
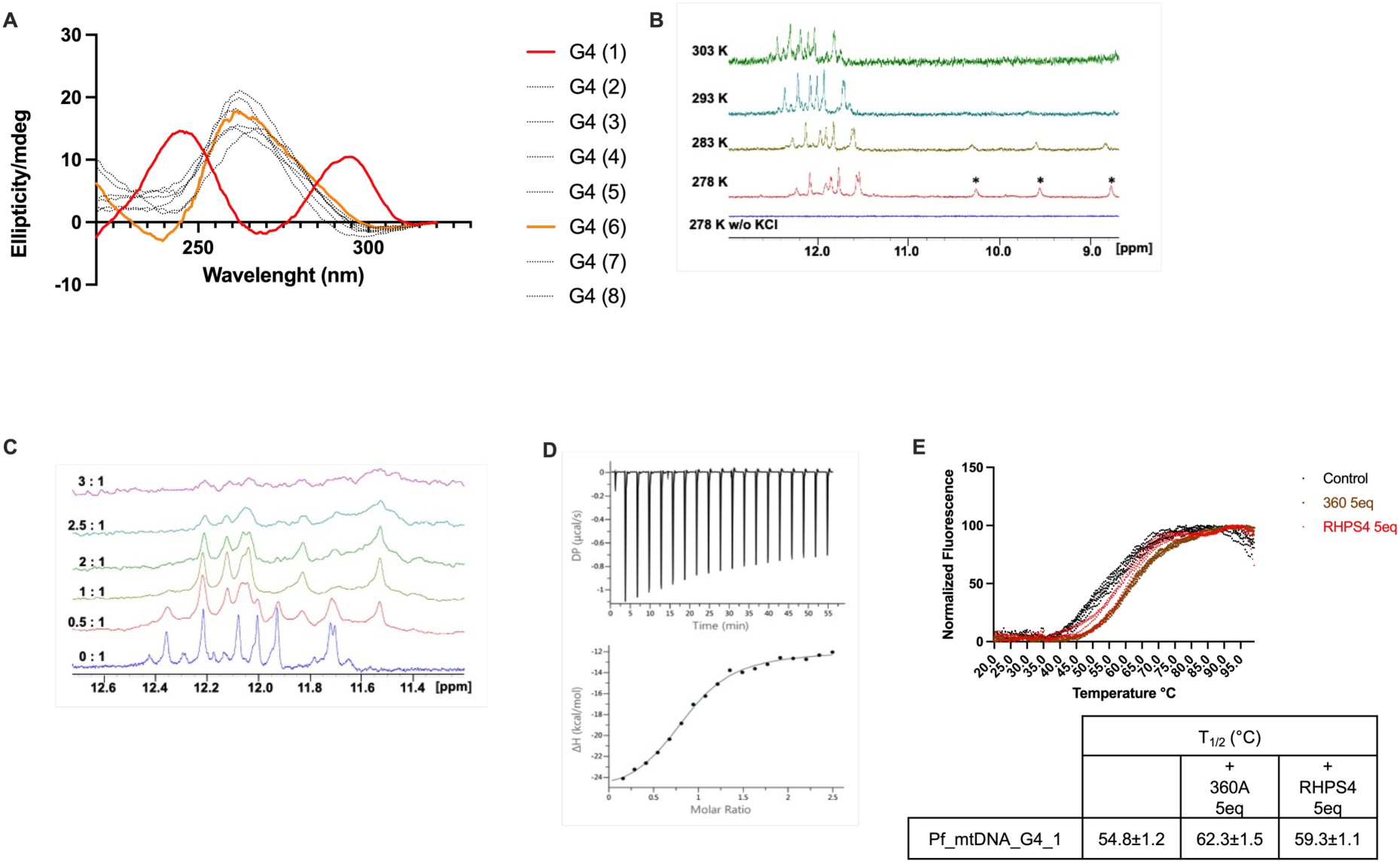
Biophysical characterization of G-quadruplex formation and RHPS4 binding to a mitochondrial DNA sequence of *Plasmodium falciparum*. **A** CD spectra result for the 8 sequences listed in Table 1 (in the same order). The only sequence for which CD results are clearly indicative of G4 formation is shown in red (Pf_mtDNA_G4_1). **B** Evaluation of G4 formation by NMR. A first spectrum was recorded at 278K for Pf_mtDNA_G4_1 (300 mM concentration) dissolved in a cacodylate buffer without KCl. ^1^H NMR spectra were then recorded at different temperatures between 278K and 303K in the presence of 100 mM KCl. **C** NMR Titration with RHPS4. NMR measurements were conducted at 600 MHz and 293 K in a buffer consisting of 20 mM cacodylate buffer, 100 mM KCl and 10 % D_2_O (NS 256). The imino proton region of the ^1^H NMR spectra of Pf_mtDNA_G4_1 was recorded with increasing concentrations of RHPS4 (12 mM stock solution in 62 % DMSO). The RHPS4/DNA ratios are indicated on the left side of the spectra. **D** ITC binding isotherm obtained at 20°C by titrating RHPS4 (200 μM concentration) to Pf_mtDNA_G4_1 (15 μM concentration). Both compounds were prepared in a buffer supplemented with 2 % DMSO. Both the primary raw data (top) and the integrated heat data (bottom) are shown. **E** Thermal stabilization of the F-Pf_mtDNA-G4-1-T oligonucleotide measured by melting FRET assays. G4 ligands were added to the reaction at 5 molar equivalents (0.4 µM) with respect to F-Pf_mtDNA-G4-1-T (0.2 µM). The T1/2 value indicates the melting temperature of the oligonucleotide in different conditions.

We then considered Nuclear Magnetic Resonance (NMR) to evaluate the capacity of the sequences to fold into G4 structures. NMR can unambiguously indicate the presence of such structures by following imino-protons that resonate at unique downfield frequencies (10-12 ppm) when involved in G-tetrads and consequently in Hoogsteen hydrogen bond networks, compared to imino protons involved in Watson-Crick base pairing that resonate between 12-14 ppm. NMR experiments were first conducted on synthetic oligonucleotides at 277 K in a lithium cacodylate buffer with 150 mM KCl. Out of the 8 considered sequences, only two (Pf_mtDNA_G4_1 and 3) gave reasonably clear and well resolved imino proton resonances in the 10-12 ppm range (Supplementary Fig. S4D). The 2D NOESY spectra recorded for sequences 1, 3 and 7 revealed the presence of several NOEs between guanine imino protons, validating the presence of G4 structures. For three sequences (Pf_mtDNA_G4_4, 5 and 6), significantly broadened signals were observed suggesting extensive exchange which prevented a clear validation of unique G4 structures. Finally, the spectra recorded for sequences Pf_mtDNA_G4_2, 7 and 8 showed a reduced number of imino signals at 10-12 ppm compared to those within the range of 12-14 ppm, suggesting a poor propensity to form G4 structures in solution.

We focused on the Pf_mtDNA_G4_1 sequence for which all our biophysical data suggested G4 folding. At 278K, the 1D spectrum showed the presence of several peaks at 11-12.5 ppm, confirming the propensity of this sequence to form G4 structures in solution. Addition of a final K^+^ concentration of 100 mM was required to obtain characteristic guanine imino protons chemical shifts involved in G-tetrad, which confirms the necessary presence of cations for G-rich oligonucleotides to fold into G4 structures (Fig. 4B). In the presence of 100 mM KCl, although the NMR spectrum did not show clear and sharp imino signals, the count of more than 8 imino protons within that range suggests the formation of at least a two-layer G4 structure. Additional peaks were found between 8 and 10 ppm, likely corresponding to hydrogen-bonded guanine amino protons within the guanine quartets (Fig. 4B). These peaks broadened with increasing temperature and disappeared above 293K, while the G4 imino signals remained observable even at 303K, reflecting relative stability of the structure.

To test if RHPS4 is able to interact with Pf_mtDNA_G4_1 sequence, titration was performed at 293K by recording 1D NMR spectra upon addition of increasing concentration of RHPS4. Effects were visible after addition of very low amounts of ligand (Fig. 4C). Notably, all proton signals significantly broadened upon the addition of RHPS4 and several shifts were observed, revealing that RHPS4 affects the G-quartets. Beyond a 1:1 ratio, no further chemical shift changes were observed indicating that the complex had fully formed. By contrast, the addition of RHPS4 to the Pf_mtDNA_G4_2 oligonucleotide which showed no evidence of G4 folding in CD experiments did not induce any chemical shift changes (Supplementary Fig. S4E).

We then applied Isothermal Titration Calorimetry (ITC) to extract thermodynamic parameters associated with the association of RHPS4 to Pf_mtDNA_G4_1 sequence. We found that the binding reaction was an exothermic process exclusively guided by a favorable enthalpy (ΔH = −14.1 kcal.mol^-1^) (Fig. 4D). The stoichiometry close to 1 suggests that the RHPS4 ligand interacts with one quadruplex in a 1:1 complex, which is fully consistent with our NMR data. Fitting the ITC data to a one-site binding model provided an association constant of (0.56± 0.08) x 10^6^ M^−1^ at 20°C which corresponds to a dissociation constant (Kd) of 1.8 ± 0.3 µM, *i.e*, in the low micromolar range.

To confirm the interaction of RHPS4 with the G4 structure formed by the Pf_mtDNA-G4-1 sequence and to evaluate its ability to stabilize this structure *in vitro*, FRET melting assays were performed (*61*). In this assay, the interaction of RHPS4 with the G4 structure is monitored through the increase in the half-melting temperature (T₁/₂) of the double-labelled G4-forming oligonucleotide F-Pf_mtDNA-G4-1-T (fluorescein (F) as the fluorescent donor and TAMRA (T) as the acceptor, attached respectively to the 5′ and 3′ ends for FRET analysis). Under 100 mM K⁺ conditions, the addition of five molar equivalents of RHPS4 relative to the oligonucleotide concentration led to a marked increase in the half-melting temperature (T₁/₂) by 4.5 °C compared with ligand-free conditions, thereby demonstrating the capacity of this compound to interact with and stabilize the structure formed by the Pf_mtDNA-G4-1 sequence. The increase in T₁/₂ observed in the presence of RHPS4 was of a similar magnitude to that obtained with the highly selective G4 ligand 360A (*61*) (ΔT₁/₂ = 7.5 °C relative to control conditions) (Fig. 4E), confirming that the observed shifts in melting temperature are consistent with the stabilization of a G4 structure formed by this oligonucleotide.

### *In vitro* RHPS4 pressure did not select for resistance

One of the major challenges that new antiplasmodial drug face upon implementation in the field is the appearance of drug-resistant *Plasmodium* parasites. To investigate the occurrence of such resistance for RHPS4, we submitted the F32-TEM parasite strain to *in vitro* discontinuous exposures of RHPS4 increasing concentrations from 90 nM up to 1.15 µM over a 17 months period. After 18 drug pressure cycles, no significant increase of the RHPS4 IC_50_ values was observed (Fig. 5A), with IC_50_ values between 64 nM and 71 nM. Moreover, there was no difference in recrudescence capacity between the selected parasite line obtained after the 18-drug pressures (F32-G4p18) and the untreated control tween strain cultured in parallel (F32-TEM), after 48 h of treatment with RHPS4 at concentrations of 250, 500 and 1000 nM (corresponding to 3, 6 and 12 times the IC_50_; *p*-value = 0.08, 0.22 and 0.43 for 250 nM, 500 nM and 1000 nM respectively; Fig. 5B). These results indicate that more than a year of RHPS4 pressure was not sufficient to select for resistance.

**Figure 5:**
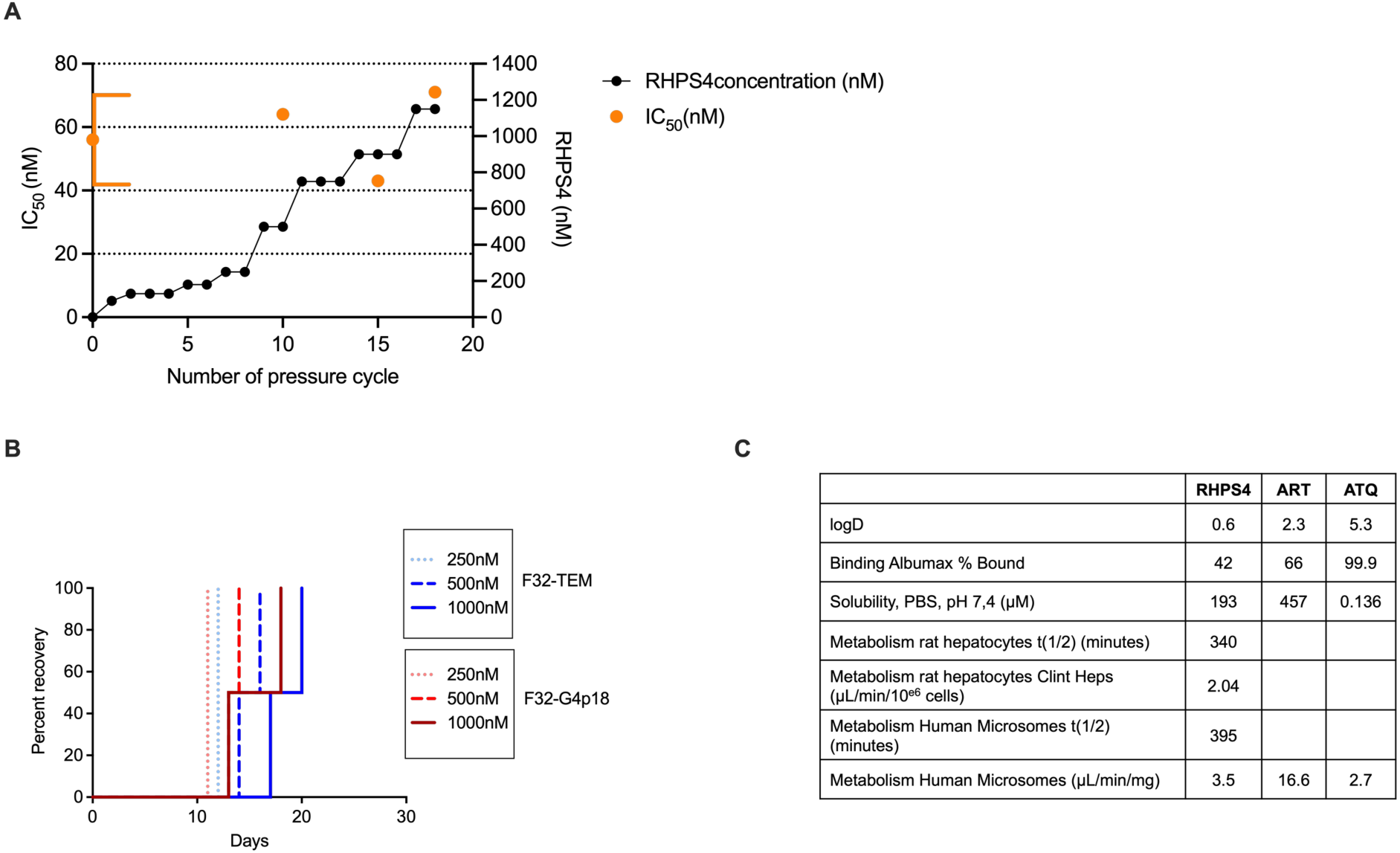
*In vitro* selection of RHPS4 pressure-adapted *Plasmodium falciparum* and and pharmacological properties. **A** Evolution of RHPS4 concentration (dark line) over the course of 18-RHPS4 pressure cycles and IC_50_ values obtained for RHPS4 (orange) at cycle 0, 10, 15 and 18. **B** Kaplan-Meier analysis of *P. falciparum* in vitro recrudescence tests comparing the ability of the RHPS4-treated strain at cycle 18 (F32-G4p18, red color gradients) and the untreated control strain cultured in parallel (F32-TEM, blue color gradients) to proliferate after treatment with RHPS4 at 250 nM, 500 nM and 1000 nM. The Mantel-Cox test was used for statistical analysis of the recrudescence data performed in two independent experiments (** = p<0.001). **C** Pharmacological properties and metabolism data of RHPS4. Artemisinin (ART) and atovaquone (ATQ) were used as references; data from MMV.

### Physico-chemical and pharmacological properties of RHPS4

Since RPHS4 activity, selectivity and original mode of action make it a promising new antiplasmodial drug, we investigated some of its physico-chemical and metabolism properties. First, the value of LogD of 0.6 at pH 7.4 indicates that RHPS4 is more hydrophilic than ART (2.3) and ATQ (5.3), (*62*)(Fig 5C). Then, with a 42% of binding to Albumax, RHPS4 displays lower interaction with human serum proteins, such as albumin, as compared to ART (66%) and ATQ (99.9%). Moreover, RHPS4 metabolization on rat hepatocytes and human microsomes is moderate and similar to ATQ, indicating a good stability (Fig. 5C).

## DISCUSSION

We have demonstrated that RHPS4, a well-characterized G4 ligand, exhibits potent antiplasmodial activity (IC₅₀ < 100 nM) against several strains of *Plasmodium falciparum*, including resistant laboratory strains and clinical isolates, with a high selectivity index (>100) relative to mammalian cells. These properties, combined with its favorable physicochemical and pharmacological profiles, position RHPS4 as a promising candidate for the development of novel antimalarial agents. A key advantage supporting its therapeutic potential is the absence of cross-resistance with front-line antimalarials such as chloroquine (CQ), atovaquone (ATQ), and artemisinin (ART). This suggests RHPS4 may be effective against multidrug-resistant malaria parasites, which are increasingly prevalent worldwide.

Stage-specific treatment assays revealed that RHPS4 exerts its antiparasitic effect primarily during the trophozoite stage (12–36 h post-invasion), a period marked by intense metabolic activity and active replication of both nuclear and mitochondrial DNA (*63*, *64*). A similar stage-dependent profile has been described for ATQ, an established inhibitor of mitochondrial electron transport (*65*, *58*). Pharmacodynamic analysis further showed that RHPS4 acts as a slow-acting compound, with a lag phase, parasite reduction ratio (PRR) and parasite clearance time (PCT) also comparable to those of ATQ. According to Sanz *et al* (*45*)., compounds with similar modes of action exert a similar killing profile. Therefore, we wondered if RHPS4 was also targeting the mitochondria. Mitochondrial staining with Mitotracker CMX-Ros confirmed a pronounced disruption of the mitochondrial membrane potential following RHPS4 treatment, a phenotype also observed with ATQ exposure (*66*, *67*). Altogether, these results strongly suggest that RHPS4 interferes with mitochondrial function. However, the lack of cross-resistance with ATQ-resistant strains suggests that RHPS4 may operates via a distinct molecular mechanism.

Molecular analyses revealed that RHPS4 significantly compromises mitochondrial DNA (mtDNA) integrity. Treatment with RHPS4 induced a 5- to 10-fold reduction in mtDNA abundance across all *P. falciparum* strains tested, correlating with a marked decrease in transcript levels of mitochondrial-encoded genes. In contrast, treatment with ART or ATQ did not affect mtDNA abundance, indicating that RHPS4 likely acts through a novel mechanism involving the disruption of mitochondrial genome maintenance and transcription. Originally characterized as a telomerase inhibitor due to its ability to bind and stabilize human telomeric G4 structures, RHPS4 has been predominantly studied in the context of telomere dysfunction (*36*, *37*, *52*). However, work by Falabella and colleagues demonstrated a preferential activity of RHPS4 toward mitochondrial nucleic acids in mammalian cells. Compared to other G4 ligands with similar *in vitro* binding and stabilization capacities, RHPS4 exhibits a unique mitochondrial localization, likely due at least in part to its net positive charge, which may promote its accumulation within mitochondria via the organelle membrane potential (*28*). Notably, *P. falciparum* harbors only a single mitochondrion per cell, in contrast to the thousands present in mammalian cells (*68*). This stark difference in mitochondria number may underlie RHPS4’s observed selectivity for the parasite over human cells.

We then wondered if the observed loss of mtDNA directly linked to RHPS4’s interaction with mitochondrial G4s. Our bioinformatic and biophysical analyses identified at least one sequence within the *P. falciparum* mitochondrial genome capable of forming a two-tetrad G4 structure. Although such structures are inherently less stable than canonical three-tetrad G4s, our data indicate that it remains stable at 30°C, suggesting functional relevance under physiological conditions. Moreover, binding studies with RHPS4 and other G4 ligands support further stabilization of this structure, consistent with a potential role in mediating RHPS4’s mitochondrial effects. Although a direct interaction between RHPS4 and mitochondrial G4s cannot be formally demonstrated, the effects of PhenDC3 and 360A (two of the most selective G4 ligands described to date) on mtDNA levels suggest a shared mechanism of action. It is worth noting that the effects of G4 ligands on *Plasmodium* mtDNA appear to depend on their overall net charge, as Pyridostatin and CX-5461, which lack a net permanent positive charge, show no significant effect on the parasite’s mtDNA. However, the marked differences in activity between RHPS4 and other positively charged G4 ligands suggest that additional factors contribute to their ability to disrupt the mitochondrial genome.

Altogether, these findings support a model in which RHPS4 selectively accumulates in parasite mitochondria and binds G4 motifs within mtDNA, leading to mtDNA depletion and collapse of mitochondrial function. This mechanism distinguishes RHPS4 from existing antimalarial drugs, including ATQ, which targets the mitochondrial cytochrome bc1 complex without altering mtDNA abundance.

The identification of Pf_mtDNA_G4_1 as a functional G4 motif provides, to our knowledge, the first direct evidence of a G-quadruplex structure in *P. falciparum* mitochondria. It suggests that mtDNA G4 structures could represent unexplored druggable targets for antimalarial strategies, particularly in the context of rising resistance to current treatments.

In conclusion, our study highlights RHPS4 as a promising antiplasmodial with a novel mechanism of action and showing no cross-resistance with other antimalarial compounds. These results open new avenues for antimalarial drug discovery targeting mitochondrial nucleic acid structures and underscore the potential of G4 ligands in combating resistant malaria.

## Materials and methods

### Drugs

Atovaquone (ATQ) and artemisinin (ART) were purchased from TCI Chemicals, chloroquine diphosphate (CQ) from Sigma Aldrich/Merck (Germany), and RHPS4 was purchased from MedChemExpress (MCE). Stock solutions were obtained by dissolving compounds in dimethyl sulphoxide (DMSO, Sigma-Aldrich) for ART, ATQ and RHPS4, and in RPMI-1640 for CQ. The percentage of DMSO in all experiments is less than or equal to 0.5%.

#### Human cell lines and culture conditions

All the cell lines used in this study were obtained from ATCC that certified their identity. DAPI staining analysis was used to confirm the absence of *Mycoplasma* contamination during the course of experiments. All culture media were provided by Gibco and were supplemented with 10% fetal bovine serum (Eurobio), 100 U/mL penicillin (Gibco), and 100 µg/mL streptomycin (Gibco). HEK, HepG2 and BJ-hTert cells were grown in Dulbecco’s Modified Eagle MediumCells in a humidified atmosphere with 5% CO_2_ at 37°C.

### Parasite strains

F32-ART is a laboratory strain, obtained after artemisinin drug pressures and carrying the M476I mutation on the *pfk13* gene responsible for its ART resistance (*38*). F32-TEM is its twin ART-sensitive strain (*39*). IPC8262 (*pfk13* C580Y, *pfcrt* M74I/N75E/K76T) is a field isolate from Cambodia resistant to both ART and CQ, provided by Dr B. Witkowski of the Pasteur Institute in Cambodia. Q120 (*pfcytb* Y268N, *pfcrt* K76T) is a field isolate from Guyana resistant to both ATQ and CQ, provided by Dr L. Musset from Pasteur Institute of Cayenne. The 3D7 lab strain is drug sensitive, Dd2 is CQ-resistant, and RF12 is both ART- and CQ-resistant (*40*).

### Parasite culture

Parasites were cultured according to Trager and Jensen (*41*) with slight modifications. All field isolates (IPC8262 and Q120) were cultured at 2% hematocrit in human red blood cells (EFS, French blood bank, France) in RPMI-1640 medium (with HEPES and L-glutamine, Dutscher, France) supplemented with 5% human serum (EFS, French blood bank, France), 0.55% Albumax II (Fisher Scientific, France), 0.4 mM hypoxanthine, 1 mM L-glutamine and 11 µg/mL gentamicin and parasites incubated with 5% O_2_, 5% CO_2_ and 90% N_2_ at 37°C. In parallel, the strains F32-TEM and F32-ART were cultivated at 2% hematocrit in human red blood cells in RPMI-1640 medium supplemented with 5% human serum, and incubated with 5% CO_2_ at 37°C in humidified atmosphere.

### Chemosensitivity assay

The SYBR Green I assay was carried out to evaluate the *in vitro* antimalarial activity of the different compounds.(*42*) D-sorbitol-synchronized ring-stage parasites were adjusted at 1% parasitemia before being exposed during 48 h at 37°C to different concentrations of the drugs in 96-well plates. Each concentration was tested in triplicate. Fluorescence was measured at 485 nm excitation / 528 nm emission in the VICTOR Nivo plate reader (Perkin Elmer, USA). The IC_50_ values were then calculated using GraphPad Prism software (USA).

### Recrudescence assay

A recrudescence assay was performed on the strain sets: F32-ART vs. F32-TEM, to compare the ability of each strain to survive and replicate after drug exposure (*43*). D-sorbitol-synchronized ring-stage parasites adjusted to 3% parasitemia and 2% hematocrit, were treated with the drug of interest in a 6-well plate. After 48 h of exposure, the drug was washed off with RPMI-1640 and the parasites were transferred to drug-free culture conditions at 10% human serum. Blood smears were made to follow recrudescence until the day that each parasite culture reached the 3% initial parasitemia, defined as the day of recrudescence. If no parasite recrudescence was observed after 30 days, the experiment was stopped and data were censored. Data analysis was performed using Kaplan-Meier survival curves. Statistical significance was determined by a log-rank (Mantel-Cox) test using GraphPad Prism software (USA).

#### Viability assay

Cells were seeded in 96-flat-wells plate at 3500 cells per well. Serial dilutions of various compounds were realized allowing same solvent concentration for each condition, and cells were treated 24 h after seeding. After 3 days, cells were fixed for 1 h at 4°C by addition of 10% trichloroacetic acid to a 3.33% final concentration, before being washed with tap water and dried overnight. Cells were stained by incubation 30 min at room temperature in a 1% acetic acid solution containing 0.057% Sulforhodamin B (SRB), then cells were washed with 1% acetic acid and dried overnight. Finally, 200 μL of a 10 mM Tris-base solution was added, plates were agitated for 1 h at room temperature, and SRB levels were measured by absorbance at 490 nm using μQuant microplate spectrophotometer (Bio-Tek Instruments). Percentages of cell viability are expressed after normalization relative to untreated cells. The IC_50_ (50% inhibitory concentration) was computed for each drug and cell lines with the GraphPad Prism software using a nonlinear regression to a four-parameter logistic curve.

### Stage-specific activity of RHPS4

To determine the time of action of RHPS4 during parasite erythrocytic development (*44*), the parasite cycle was subdivided into five distinct time windows: 0-12 h, 12-24 h, 24-36 h, 36-48 h, and 48-60 h to cover one and a quarter parasite cycle. The complete cycle condition (0-48 h) was performed as a control, and an extended condition (0-60 h) was added to assess the effect of the molecule on parasite reinvasion. Synchronized ring-stage F32-TEM parasites (0-6 h post-invasion, synchronized by percoll/sorbitol treatment) adjusted to 0.5% parasitemia and 2% hematocrit were divided into 24-well plates. Wells were assigned to each time slot and treated with three concentrations of the test molecule corresponding to the value of IC_50_, 2 × IC_50_ and 5 × IC_50_, while an untreated well served as a control. Each treatment lasted 12 h and after washing to eliminate the molecule, the parasites were cultured in drug-free medium until the end of the last treatment. At the end of the 60 h, blood smears were prepared. In addition, SYBR Green labelling was performed for each condition and the percentage inhibition was determined based on the amount of labelled DNA, which reflects parasite proliferation.

#### *In vitro* Parasite Reduction Ratio Assay (PRR)

The aim of this test was to assess the parasite killing rate (PRR) in response to treatment. It was carried out in accordance with the method described by Sanz *et al* (*45*) with slight modifications. For this test, highly synchronized ring-stage F32-TEM parasites (0-4 h post-invasion, synchronized by percoll/sorbitol treatment) adjusted to 0.5% parasitemia and 2% hematocrit (corresponding to 1 x 10^6^ parasites per mL) were exposed in 6-well plates to the compound to be tested at a concentration corresponding to 10 times its IC_50_. Every 24 h for up to 120 h, culture was washed with RPMI-1640 and an aliquot transferred in a 96-well plate in which a serial dilution (1:3) was performed in uninfected RBC at 2% hematocrit. In the 6-well plate, parasites were treated again with the compound to be tested and incubated until the next time point 24 h later. The 96-well plates were maintained in culture for 3 weeks before SYBR Green labelling to determine the number of wells in which parasites proliferated. The number of parasites alive at each time point is determined using the following formula: X^n-1^; Where X represents the serial dilution factor (here = 3) and n is the number of wells in which parasites grow. The log of (viable parasites + 1) is plotted *versus* time and used to determine the Log(PRR), which is difference in the number of viable parasites at the end of the lag phase and 48 h later. The lag phase is defined as the time needed to reach the maximum speed of action. 99.9% parasite clearance time (PCT) is the time needed to decrease parasite number by 3 logs.

#### Mitochondrial labelling with the MitoTracker Red CMXRos

The Mitotracker dye was prepared according to the manufacturer’s recommendations (Invitrogen, USA) in DMSO at 1 mM and stored at −20°C until use. A 1 µM working solution in pre-warmed RPMI-1640 was prepared directly from the stock solution just prior to parasites staining in culture. F32-TEM parasites synchronized by percoll/sorbitol treatment at the trophozoite stage (24-28 h post-invasion) were exposed for 6 h to two different conditions: a control condition (untreated) and a condition treated with RHPS4 at 180 nM (2 × IC_50_). At the end of the treatment, the drug was washed off with RPMI-1640. The parasites were adjusted to 2.5% hematocrit, and 500 nM of CMXRos MitoTracker Red were added. After one hour of incubation in the dark, the parasites were washed three times with RPMI-1640, and then 2 µg/mL of DAPI was added. The parasites were then incubated for 10 minutes in the dark, after which they were washed with RPMI-1640 and PBS 1×. They were then suspended in RPMI-1640 to prepare blood smears. These were fixed in 100% ethanol for 10 minutes and then mounted in VectaShield and sealed with varnish. Images were taken with a Zeiss Elyra 7 super-resolution microscope at 63× magnification. The MitoTracker Red CMXRos excitation wavelength is 579 nm and the emission wavelength is 599 nm. For DAPI, the excitation wavelength is 350 nm and the emission wavelength is 465 nm. Fluorescence intensities were measured using ImageJ software.

### In vitro potentiation test

The effect of ATQ on RHPS4 activity in F32-TEM parasites was determined by potentiation experiments, as described by Benoit-Vical *et al*. (*46*) D-sorbitol-synchronized ring-stage parasites adjusted to 1% parasitemia and 2% hematocrit, were exposed to different concentrations of the molecules alone (RHPS4 and ATQ) or to combinations of the two molecules, in a 96-well plate. After 72 h incubation, cells were washed with PBS 1× three times and labeled with SYBR Green I 1× for 1 h. IC_50_ values for the single molecule and the combination are then calculated using GraphPad Prism software (USA). Isobolograms are constructed by plotting a pair of fractional IC_50_ for each combination of RHPS4 and ATQ. Fractional IC_50_ for each molecule were calculated by dividing the IC_50_ value for the combination by the IC_50_ value for the molecule alone. The experiment was run three times with 3 technical replicates.

### Relative quantification of mitochondrial DNA (qPCR), and RNA (qRT-PCR)

Total genomic DNA was extracted directly from *P. falciparum* infected RBC as described in the manufacturer’s user manual of the High Pure PCR Template Preparation Kit (Roche). To measure the relative abundance of mitochondrial DNA, we designed primers for mitochondrial sequences of interest (Supplementary Table 1). Real-time PCR (qPCR) using SsoFast Eva Green Supermix (Bio-Rad), 10 ng/reaction DNA and 300 nM primers in 10 μL of final reaction volume, was performed in a thermal cycler (Bio-Rad) with the following parameters: 95°C for 2 min; 39 cycles of 95°C for 10 s, 60°C for 10 s followed by a melting curve from 65 to 95°C. The relative abundance of mitochondrial DNA was calculated by the ΔΔCt method using the nuclear gene PF3D7_1371000, which encodes ribosomal RNA 18S, for normalization. To quantify the relative abundance of mRNA or rRNA by real-time quantitative polymerase chain reaction (qRT-PCR), total RNA was isolated from parasite pellets (after RBC lysis by saponin treatment) using the RNeasy Mini kit (Qiagen, Germantown, MD). Next, cDNA was synthesized using the iTaq Universal SYBR Green One-Step Kit (Bio-Rad) with specific primers (Supplementary Table 1), 20 ng/reaction and 300 nM primers in 10 μL of final reaction volume in a thermal cycler (Bio-Rad) with the following parameters: 50°C for 10 min, followed by 95°C for 1 min and 39 cycles at 95°C for 10 seconds and 60°C for 30 seconds, followed by a melting curve from 65 to 95°C. The relative abundance of RNA was calculated according to the ΔΔCt method using the nuclear gene PF3D7_1121300, which encodes for the protein tyrosine kinase-like TLK2, for normalization.

#### Spectroscopy

For these studies, oligonucleotides were purchased from Sigma and used without further purification. Stock solutions were prepared at 100 μM strand concentration in pure water. All oligonucleotides were annealed in a lithium cacodylate buffer supplemented with 100 mM KCl, kept at 95°C for 5 min and slowly cooled to room temperature before measurements. Circular Dichroism (CD), differential absorbance spectra (IDS and TDS) and iso-FRET experiments were performed as previously described (*47–49*). The experiments were performed at 25°C in a lithium cacodylate buffer in the presence of 100 mM potassium ions.

FRET-melting experiments were carried out in 96-well plates on a CFX96 real-time PCR equipment (Bio Rad). After an initial stabilisation at 20 °C for 10 min, the temperature was increased by a 0.5 °C step every minute until 99 °C. The labelled oligonucleotide was dissolved in stock pH 7.2 buffer (10 mM lithium cacodylate, 90 mM lithium chloride) to 100 µM stock concentration and stored at −20 °C. The melting experiments were performed at a final 0.2 µM strand concentration of oligonucleotide with 5molar equivalents of ligands relative to oligonucleotide concentration. The measurement buffer consisted in 10 mM lithium cacodylate pH 7.2 buffer, 100 mM potassium chloride. The F-Pf_mtDNA-G4-1-T (5’-FAM-**TGGATGGTGTTGGCTGGGC**-TAMRA-3’) was purchased from EUROGENTEC.

#### NMR experiments

Purified oligonucleotides were purchased from Eurogentec. Unless specified, powders were resuspended into a 20 mM lithium cacodylate buffer with 100 mM KCl at pH 6.8. The oligonucleotides were heated to 95°C for 5 min. and subsequently cooled down slowly to room temperature. For NMR titrations, RHPS4 powder was resuspended into deuterated DMSO to reach a final concentration of 12.5 mM in 62% DMSO. NMR experiments were recorded using Bruker 600 and 700 MHz spectrometers equipped with a cryoprobe.

#### Isothermal Titration calorimetry experiments

Isothermal Titration calorimetry experiments (ITC) were performed at 10 and 20°C using a PEAT-ITC instrument (Malvern Panalytical). Both ligand and DNA solutions were diluted in a cacodylate buffer with 100 mM KCl and 2% DMSO to allow correct ligand solubility. Titration cell was filled with a solution of 20-30 μM oligonucleotide while the syringe was loaded with a RHPS4 solution of 200-300 μM. Experiments consisted of a first injection followed by 18 injections of 4 s with a 180 s interval between the injections. Control dilution experiments were performed by injecting ligand solution into the buffer alone placed in the measurement cell. The ITC titrations were processed using the MicroCal Origin software.

#### Pf-LDH test

*P. falciparum 3D7* (From BEI resource) lactate dehydrogenase (Pf-LDH) growth inhibition assay was carried out as described in (*50*) with minor modification in the culture state of inoculum (10-15% parasitaemia with ≥80% rings).

Data were normalized to percent growth inhibition with respect to positive (0.2% DMSO as 0% inhibition) and negative (mixture of 100mM Chloroquine and 100mM Atovaquone as 100% inhibition) controls.

#### Study of metabolic stability in rat hepatocytes

A solution of the test compound in Krebs-Henseliet buffer solution (1 µM) was incubated in pooled rat hepatocytes (1×10^6^cells/mL) for 0,15,30,45,60, 75 and 90minutes at 37°C (5% CO_2_, 95% relative humidity). The reaction was terminated with the addition of ice-cold acetonitrile containing system suitability standard at designated time points. The sample was centrifuged (4200 rpm) for 20 minutes at 20 °C and the supernatant was half diluted in water and then analyzed by means of LC-MS/MS. % Parent compound remaining, half-life (T_1/2_) and clearance (CL_int,app_) were calculated using standard methodology. The experiment was carried out in duplicate. Diltiazem, 7-ethoxy coumarin, propranolol and midazolam were used as reference standards.

#### Metabolic stability study using human liver microsomes

A solution of the test compounds in phosphate buffer solution (1 µM) was incubated in pooled human liver microsomes (0.5 mg/mL) for 0, 5, 20, 30, 45 and 60 minutes at 37°C in the presence and absence of NADPH regeneration system (NRS). The reaction was terminated with the addition of ice-cold acetonitrile containing system suitability standard at designated time points. The sample was centrifuged (4200 rpm) for 20 minutes at 20°C and the supernatant was half diluted in water and then analyzed by means of LC-MS/MS.% Parent compound remaining, half-life (T_1/2_) and clearance (CL_int,app_) were calculated using standard methodology. The experiment was carried out in duplicate. Verapamil, diltiazem, phenacetin and imipramine were used as reference standards.

#### Selection of RHPS4-resistant strain *in vitro*

RHPS4 resistance was selected by sequential exposure of F32-TEM parasites to several cycle of drug pressure according to the method described by Witkowski *et al*, with slight modifications (*51*). Briefly, D-sorbitol-synchronized ring-stage parasites adjusted to 3% parasitemia and 3% hematocrit were exposed to sequential increasing concentrations of RHPS4 (from 90 nM to 1.15 µM). After 48 h of incubation, the drug was washed off once with RPMI-1640 and the parasites were transferred to drug-free culture conditions at 10% human serum. Blood smears were made to follow recrudescence until the day that parasite culture reached its initial parasitemia, defined as the day of recrudescence. Parasite culture was then continued under normal conditions (5% human serum) until the next drug pressure cycle. In parallel, a flask of F32-TEM was maintained in continuous culture under the same conditions without any drug pressure as control. For selection experiments, parasites were grown with 5% O_2_, 5% CO_2_ and 90% N_2_ at 37°C.

#### Statistical analyses

Unless otherwise specified, all results are based on at least three independent experiments. Statistical analyses were performed using GraphPad Prism software. In all figures, statistically significant differences between specified conditions are indicated by asterisks (\**p* < 0.05; \*\**p* < 0.01; \*\*\**p* < 0.001; \*\*\*\**p* < 0.0001). NS denotes a non-significant difference.

## Acknowledgments

The authors acknowledge Dr Benoît Witkowski (Institut Pasteur du Cambodge, Cambodia) for sharing the Cambodian isolate IPC8262, and Dr L. Musset from Pasteur Institute of Cayenne for sharing the Q120 isolate. We would like to thank Medicines for Malaria Venture (MMV), in particular Didier Leroy and Anna Adam Sulakova, for their technical and scientific contributions, as well as the various constructive discussions on our work. We acknowledge TRI-IPBS imaging facilities (France-BioImaging ANR-24-INBS-0005 FBI BIOGEN). This work was supported by the Agence Nationale de la Recherche (ANR Project MAG4 grant ANR-22-CE18), the Fondation pour la Recherche Médicale (≪Équipe FRM≫ Grant number EQU202103012596, France), the CNRS (Centre National de la Recherche Scientifique, France) and the INSERM (Institut National de la Santé et de la Recherche Médicale, France). YL was founded by Chinese Scholarship Council fellowship 201906340018 and the fondation de l’Ecole Polytechnique - « Pathogens » project. MS was founded by a graduate fellowship program from the University Paul Sabatier of Toulouse and Région Occitanie.

## Author contributions

M.S., L.P., T.R. and F.N. carried out cellular and molecular experiments on *P. falciparum.* Y.L. performed biophysical analyses. V.G. performed NMR analyses. L.P., J-M.A., F.B-V. and D.G. designed the experiments. M.S., L.P., F.N., J-M.A., V.G., S.B., J-L.M., F.B-V. and D.G. co-wrote the manuscript.

## Supplementary Figures and tables

**Supplementary Figure S1:**
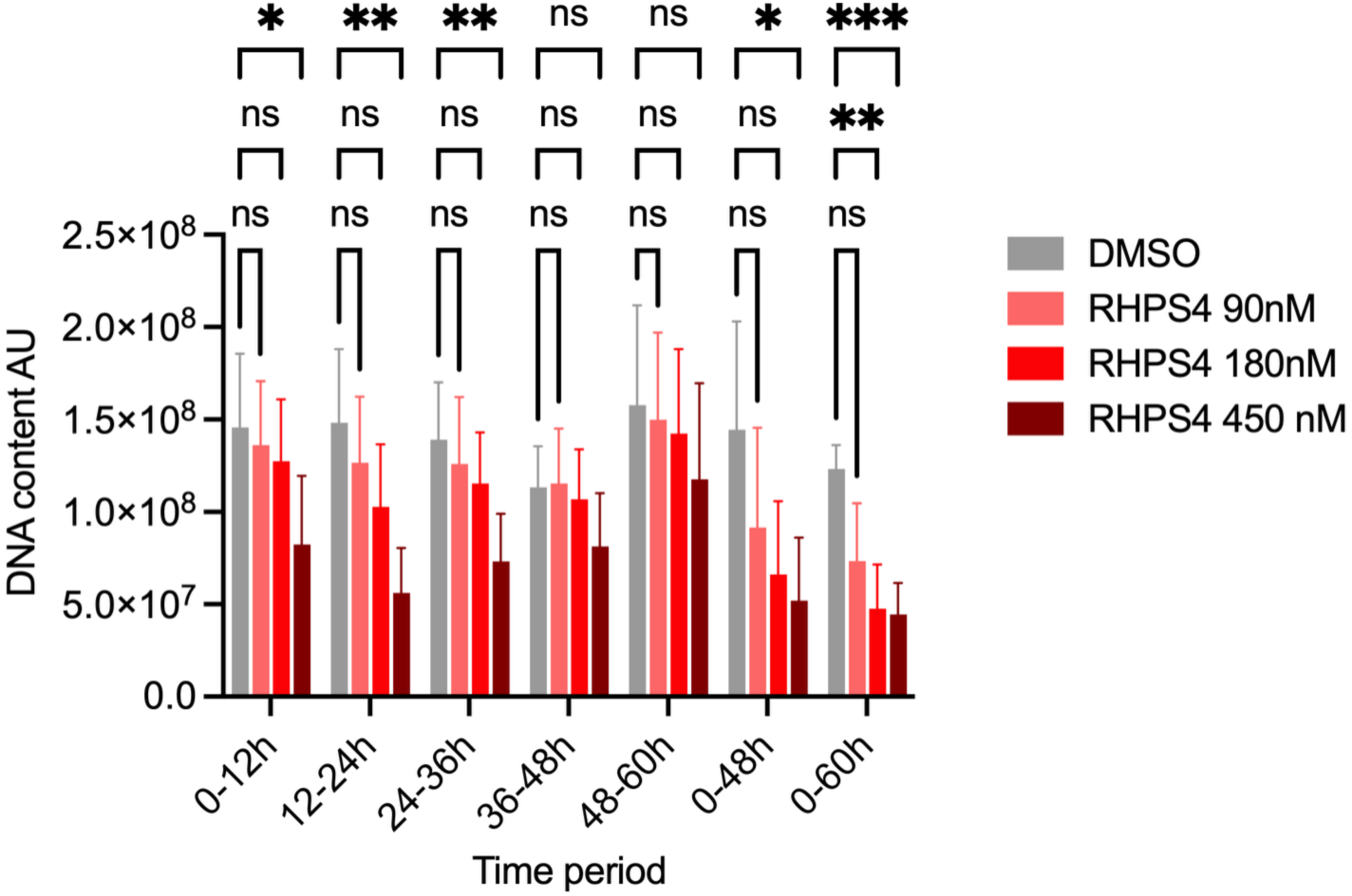
Stage specific activity of RHPS4 evaluated by DNA quantification. The F32-TEM strain was treated with RHPS4 at the 48 h IC₅₀ value, 2 × IC₅₀, or 5 × IC₅₀, initiated at successive 12-h intervals during five successive 12-h periods from a highly-synchronized ring population. Following each pulse, parasites were washed and cultured drug free until 60 h. In parallel to the smear monitoring (cf Fig. 1e), the DNA content was here determined using SYBR Green labelling. Error bars indicate mean ± SD (n = 3 independent experiments).

**Supplementary Figure S2:**
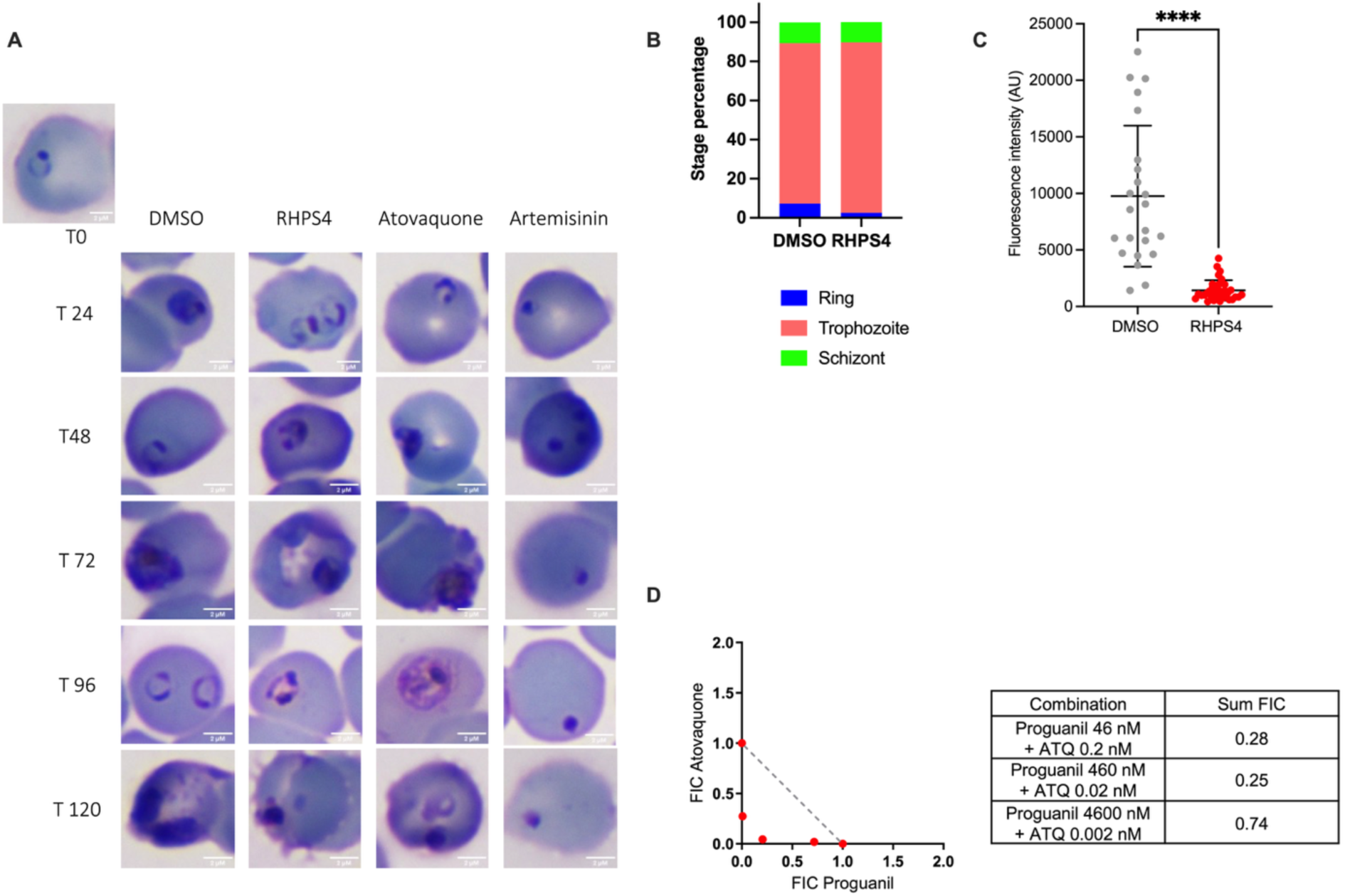
Effects of RHPS4 on *Plasmodium falciparum* morphology, mitochondrial activity, and drug interaction compared with reference compounds. **A** Giemsa smear showing the morphology over time of *P. falciparum* from a ᴅ-Sorbitol synchronized population after exposure to RHPS4 (900 nM), atovaquone (50 nM), and artemisinin (200 nM). **B** Percentage of each parasite stages (ring, trophozoite and schizont) counted by two independent experienced microscopists on Giemsa-stained blood smears. Morphologically, ring-stage parasites were defined as parasites with a ring shape, trophozoite stages as parasites with full cytoplasm, and schizonts as parasites with punctate nuclear coloration of the developing merozoites. **C** Mitochondrial fluorescence was quantified with ImageJ software by measuring the mean MitoTracker intensity per infected RBC. Bars show mean ± SD n=24 infected RBC for DMSO and n=34 infected RBC for RHPS4; *p* values were calculated with a two tailed Student’s t test. **D** Isobologram and sum of fractional IC_50_ (FIC) of interaction between atovaquone and proguanil (control combination) with *P. falciparum* F32-TEM strain.

**Supplementary Figure S3:**
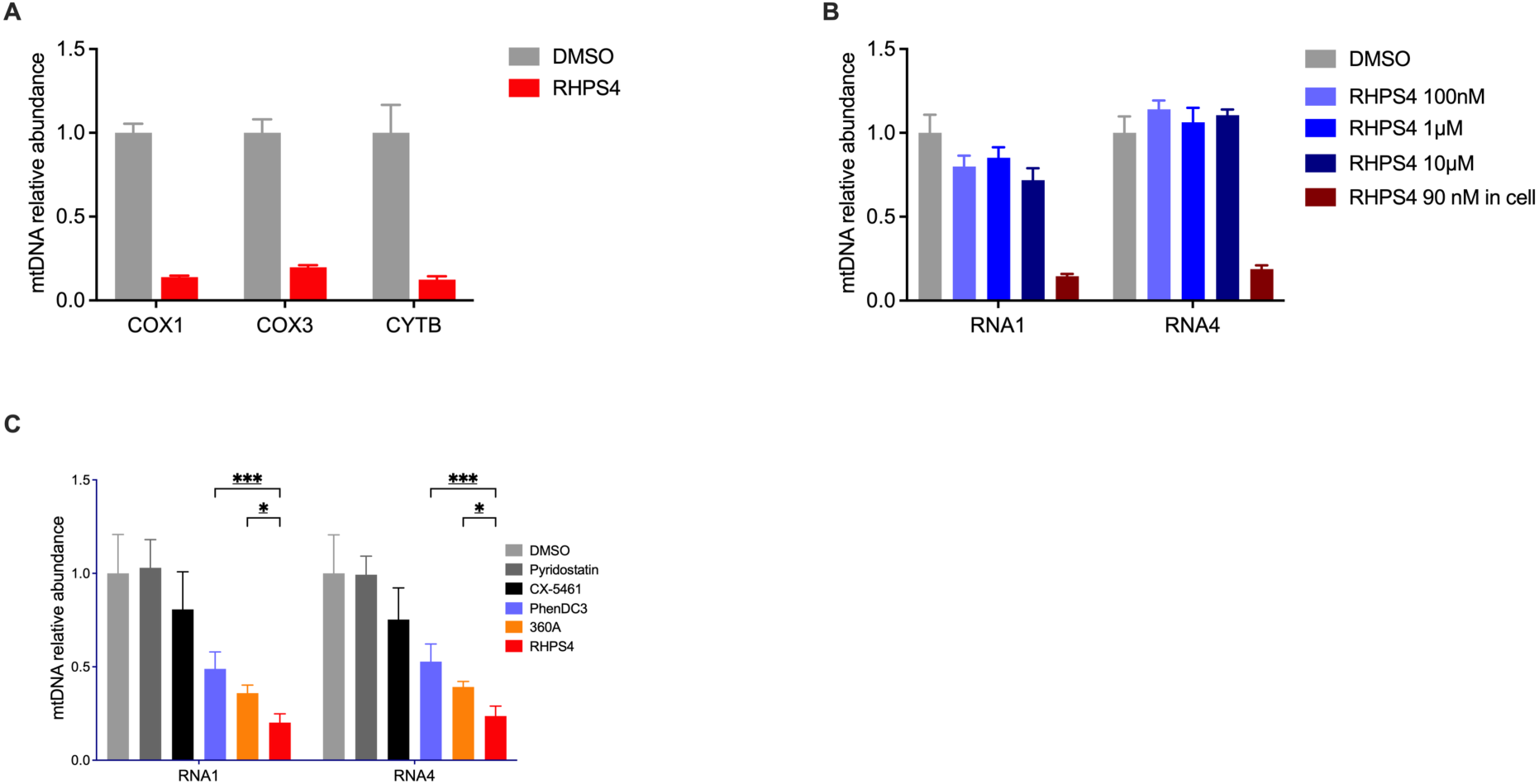
Effects of RHPS4 and G4 ligands on mitochondrial DNA abundance. **A** Relative abundance of mtDNA sequences measured by qPCR after a 48 h exposure to 90 nM RHPS4 of trophozoite stage from F32-TEM (synchronized by sorbitol), n=1. The mtDNA abundance was normalized to the nuclear sequence PF3D7_1371000 coding for ribosomal RNA 18S.**B** Assessment of RHPS4 interference with the PCR reaction. The relative abundance of two mtDNA sequences extracted from untreated F32-TEM trophozoites were measured by qPCR in presence of different concentration of RHPS4. As a control, relative quantification of mtDNA sequences from RHPS4-treated parasites was performed in parallel, using the same PCR conditions. **C** Differential effects of RHPS4 on mitochondrial DNA abundance compared to two other G4 ligands, 360A and PhenDC3. Relative abundance of mtDNA sequences measured by qPCR after 48 h of exposure to different G4 ligands at their respective IC₅₀ concentrations. mtDNA abundance was normalized to the nuclear gene PF3D7_1371000, which encodes 18S ribosomal RNA. Bars show mean ± SD from n = 3 independent experiments.

**Supplementary Figure S4:**
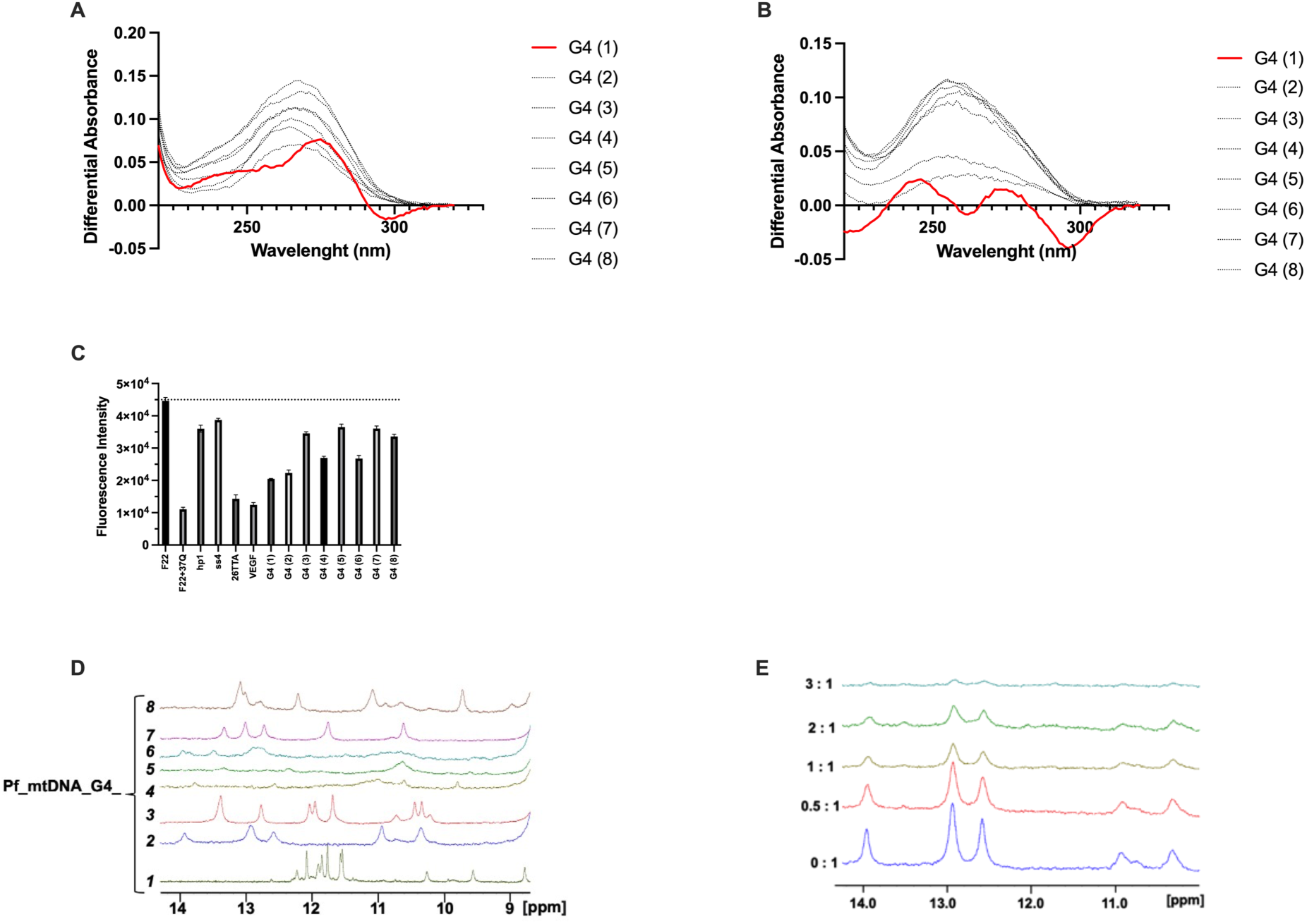
Biophysical characterization of G-quadruplex formation of mitochondrial sequences from *Plasmodium falciparum*. **A** TDS, **B** IDS and **C** iso-FRET results for the 8 sequences listed in Table 1 (in the same order). The only sequence for which CD/TDS/IDS results are clearly indicative of G4 formation is shown in red (Pf_mtDNA_G4_1). **D** Superimposition of the ^1^H NMR spectra recorded at 277 K for the eight considered sequences in a cacodylate buffer with 150 mM KCl at pH 6.5. **E** NMR Titration of Pf_mtDNA_G4_2 with RHPS4. NMR measurements were conducted at 600 MHz and 278 K in 20 mM cacodylate buffer, 100 mM KCl and 10 % D_2_O (256 scans). The imino proton regions of the ^1^H NMR spectra of Pf_mtDNA_G4_1 was recorded with increasing concentrations of RHPS4 (12 mM stock solution in 62 % DMSO). The RHPS4/DNA ratios are indicated on the left side of the spectra.

**Supplementary table 1:**
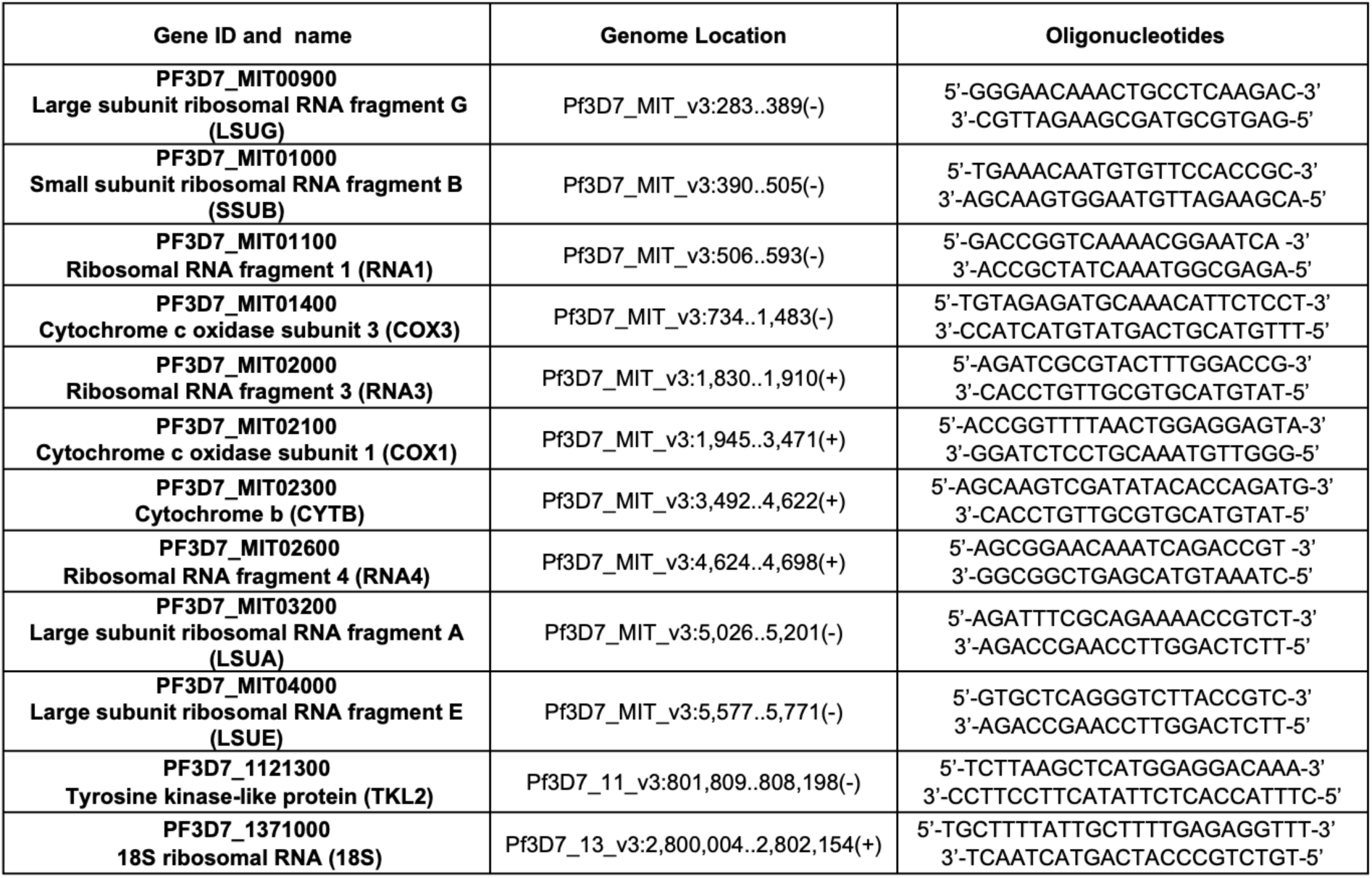
The gene name and ID and genome location, of the oligonucleotides used for qPCR, and RT-qPCR as indicated in the PlasmoDB database.

**Supplementary table 2:** The gene name and ID, genome location of *P. falciparum* mitochondrial DNA sequences susceptible to G4 structuring as indicated in the PlasmoDB database.

| Sequence name | Gene ID and name | Genome Location |
| --- | --- | --- |
| **Pf_mtDNA_G4_1** | **PF3D7_MIT00300**<br>Ribosomal RNA fragment RNA9 | Pf3D7_MIT_v3:72..125(+) |
| **Pf_mtDNA_G4_2** | **PF3D7_MIT01500**<br>Large subunit ribosomal RNA fragment F | Pf3D7_MIT_v3:1,515..1,578(+) |
| **Pf_mtDNA_G4_3** | **PF3D7_MIT02100**<br>Cytochrome c oxidase subunit 1 | Pf3D7_MIT_v3:1,945..3,471(+) |
| **Pf_mtDNA_G4_4** | **PF3D7_MIT02100**<br>Cytochrome c oxidase subunit 1 | Pf3D7_MIT_v3:1,945..3,471(+) |
| **Pf_mtDNA_G4_5** | **PF3D7_MIT02100**<br>Cytochrome c oxidase subunit 1 | Pf3D7_MIT_v3:1,945..3,471(+) |
| **Pf_mtDNA_G4_6** | **PF3D7_MIT04000**<br>Large subunit ribosomal RNA fragment E | Pf3D7_MIT_v3:5,577..5,771(-) |
| **Pf_mtDNA_G4_7** | **PF3D7_MIT04000**<br>Large subunit ribosomal RNA fragment E | Pf3D7_MIT_v3:5,577..5,771(-) |
| **Pf_mtDNA_G4_8** | **PF3D7_MIT04200**<br>Ribosomal RNA fragment RNA8 | Pf3D7_MIT_v3:5,861..5,954(-) |

